# Decoding the Information Structure Underlying the Neural Representation of Concepts

**DOI:** 10.1101/2021.03.16.435524

**Authors:** Leonardo Fernandino, Jia-Qing Tong, Lisa L. Conant, Colin J. Humphries, Jeffrey R. Binder

## Abstract

The nature of the representational code underlying conceptual knowledge remains a major unsolved problem in cognitive neuroscience. We assessed the extent to which different representational systems contribute to the instantiation of lexical concepts in high-level, heteromodal cortical areas previously associated with semantic cognition. We found that lexical semantic information can be reliably decoded from a wide range of heteromodal cortical areas in frontal, parietal, and temporal cortex. In most of these areas, we found a striking advantage for experience-based representational structures (i.e., encoding information about sensory-motor, affective, and other features of phenomenal experience), with little evidence for independent taxonomic or distributional organization. These results were found independently for object and event concepts. Our findings indicate that concept representations in heteromodal cortex are based, at least in part, on experiential information. They also reveal that, in most heteromodal areas, event concepts have more heterogeneous representations (i.e., they are more easily decodable) than object concepts, and that other areas beyond the traditional “semantic hubs” contribute to semantic cognition, particularly the posterior cingulate gyrus and the precuneus.

## Introduction

The capacity for conceptual knowledge is arguably one of the most defining properties of human cognition, yet it is still unclear how concepts are represented in the brain. Recent developments in functional neuroimaging and computational linguistics have sparked renewed interest in elucidating the information structures and neural circuits underlying concept representation (1–5). Attempts to characterize the representational code for concepts typically involve information structures based on three qualitatively distinct types of information, namely, taxonomic, experiential, and distributional information. As the term implies, a taxonomic information system relies on category membership and inter-category relations. Our tendency to organize objects, events and experiences into discrete categories has led most authors – dating back at least to Plato (6) – to take taxonomic structure as the central property of conceptual knowledge (7). The taxonomy for concepts is traditionally seen as a hierarchically structured network, with basic level categories (e.g. “apple”, “orange”) grouped into superordinate categories (e.g., “fruit”, “food”), and subdivided into subordinate categories (e.g., “Gala apple”, “tangerine”) (8). A prominent account in cognitive science maintains that such categories are represented in the mind/brain as purely symbolic entities, whose semantic content and usefulness derive primarily from how they relate to each other (9, 10). Such representations are seen as qualitatively distinct from the sensory-motor processes through which we interact with the world, much like the distinction between software and hardware in digital computers.

An “experiential” representational system, on the other hand, encodes information about the experiences that led to the formation of particular concepts. It is motivated by a view, often referred to as embodied, grounded, or situated semantics, in which concepts arise primarily from generalization over particular experiences, as information originating from the various modality-specific systems (e.g., visual, auditory, tactile, motor, affective, etc.) is combined and re-encoded into progressively more schematic representations that are stored in memory. Since, in this view, there is a degree of continuity between conceptual and modality-specific systems, concept representations are thought to reflect the structure of the perceptual, affective and motor processes involved in those experiences (11–14).

Finally, “distributional” information pertains to statistical patterns of co-occurrence between lexical concepts (i.e., concepts that are widely shared within a population and denoted by a single word) in natural language usage. As is now widely appreciated, these co-occurrence patterns encode a substantial amount of information about word meaning (15–17). Although word co-occurrence patterns primarily encode contextual associations, such as those connecting the words “cow”, “barn”, and “farmer”, semantic similarity information is indirectly encoded, since words with similar meanings tend to appear in similar contexts (e.g., “cow” and “horse”, “pencil” and “pen”). This has led some authors to propose that concepts may be represented in the brain, at least in part, in terms of distributional information (15, 18).

Whether, and to what extent, each of these types of information plays a role in the neural representation of conceptual knowledge is a topic of intense research and debate. A large body of evidence has emerged from behavioral studies, functional neuroimaging experiments, and neuropsychological assessment of patients with semantic deficits, with results typically interpreted in terms of taxonomic (19–24), experiential (13, 25–34), or distributional (2, 3, 5, 35, 36) accounts. However, the extent to which each of these representational systems plays a role in the neural representation of conceptual knowledge remains controversial (23, 37, 38), in part because their representations of common lexical concepts are strongly inter-correlated: patterns of word co-occurrence in natural language are driven in part by taxonomic and experiential similarities between the concepts to which they refer, and the taxonomy of natural categories is systematically related to the experiential attributes of the exemplars (39–41). Consequently, the empirical evidence currently available is unable to discriminate between these representational systems.

Several computational models of concept representation have been proposed based on these structures. While earlier models relied heavily on hierarchical taxonomic structure (42, 43), more recent proposals have emphasized the role of experiential and/or distributional information (34, 44–46). The model by Chen and colleagues, for example, showed that graded taxonomic structure can emerge from the statistical coherent covariation found across experiences and exemplars without explicitly coding such taxonomic information per se. Other models propose that concepts may be formed through the combination of experiential and distributional information (44, 46), suggesting a dual representational code akin to Paivio’s dual coding theory (47).

We investigated the relative contribution of each representational system by deriving quantitative predictions from each system for the similarity structure of a large set of concepts and then using representational similarity analysis (RSA) with high-resolution functional MRI (fMRI) to evaluate those predictions. Unlike the more typical cognitive subtraction technique, RSA focuses on the information structure of the pattern of neural responses to a set of stimuli (48). For a given stimulus set (e.g., words), RSA assesses how well the representational similarity structure predicted by a model matches the neural similarity structure observed from fMRI activation patterns (Figure 1). This allowed us to directly compare, in quantitative terms, predictions derived from the three representational systems.

**Figure 1:**
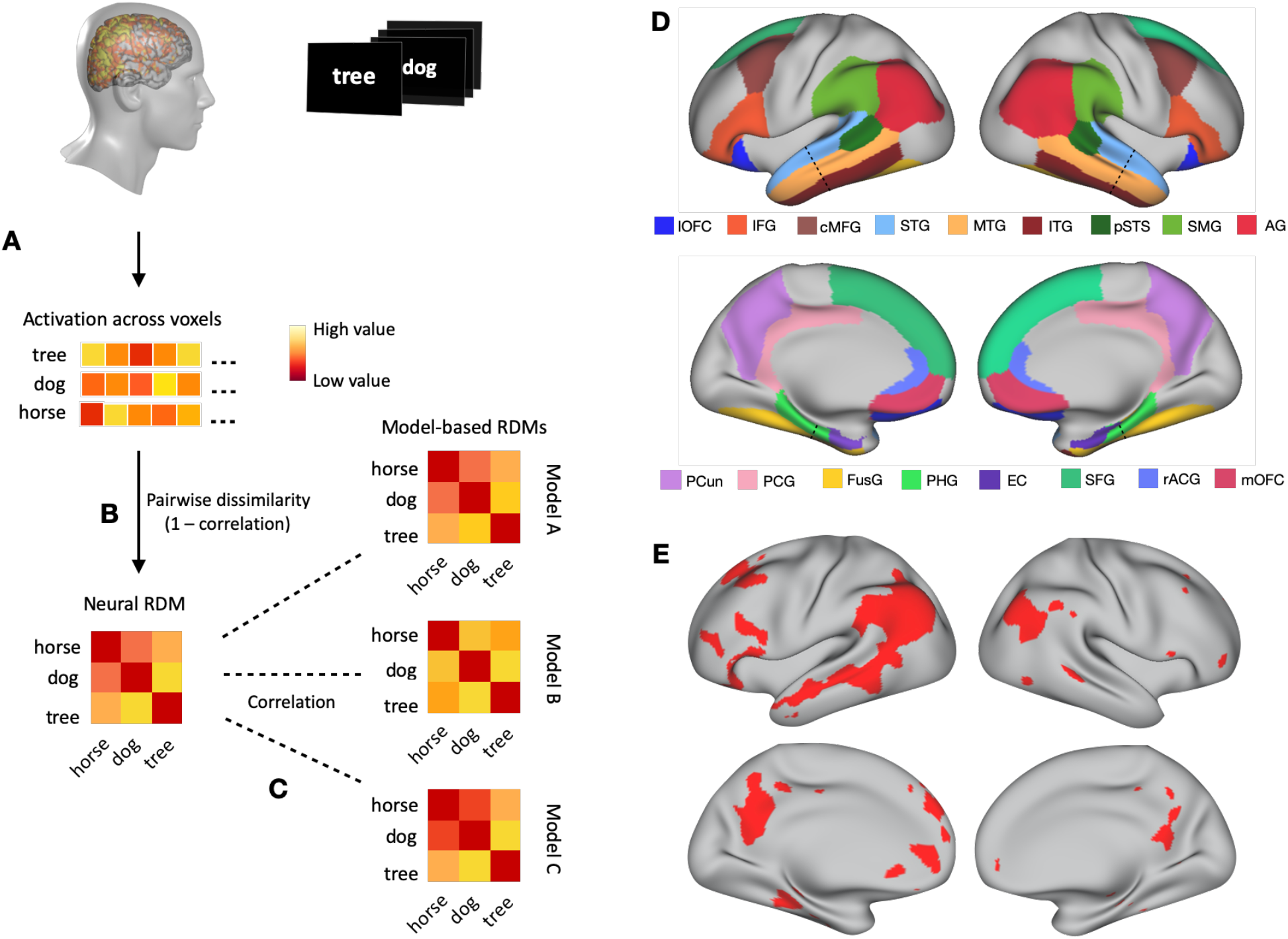
Representational similarity analysis. A. An fMRI activation map was generated for each concept presented in the study, and the activation across voxels was reshaped as a vector. B. The neural representational dissimilarity matrix (RDM) for the stimulus set was generated by computing the dissimilarity between these vectors (1 – correlation) for every pair of concepts. C. A model-based RDM was computed from each model, and the similarity between each model’s RDM and the neural RDM was evaluated via Spearman correlation. D. Anatomically defined ROIs. The dashed line indicates the boundary where temporal lobe ROIs were split into anterior and posterior portions (see main text for acronyms). E. Cortical areas included in the functionally defined semantic network ROI (49).

## Results

In two experiments, participants made familiarity judgments on a large number of lexical concepts, selected from a broad range of taxonomic categories, while undergoing fMRI (see Materials and Methods section for details). Written nouns were presented, one at a time, on a computer screen, and participants rated each one according to how often they encountered the corresponding entity or event in their daily lives, on a scale from 1 (“rarely or never”) to 3 (“often”). RSAs were conducted for a set of cortical areas previously associated with concept representation (49–51): angular gyrus (AG), supramarginal gyrus (SMG), temporal pole (TP), anterior superior temporal gyrus (aSTG), posterior superior temporal gyrus (pSTG), anterior middle temporal gyrus (aMTG), posterior middle temporal gyrus (pMTG), anterior inferior temporal gyrus (aITG), posterior inferior temporal gyrus (pITG), posterior superior temporal sulcus (pSTS), anterior parahippocampal gyrus (aPHG), posterior parahippocampal gyrus (pPHG), entorhinal cortex (EC), inferior frontal gyrus (IFG), caudal middle frontal gyrus (cMFG), superior frontal gyrus (SFG), precuneus (PCun), posterior cingulate gyrus (PCG), rostral anterior cingulate gyrus (rACG), medial orbitofrontal cortex (mOFC), lateral orbitofrontal cortex (lOFC) and fusiform gyrus (FusG). These regions were anatomically defined following the Desikan-Killiany probabilistic parcellation map (52), with the border between anterior and posterior temporal lobe areas determined according to a plane perpendicular to the lobe’s main axis (53). We also conducted RSA for a distributed, functionally defined region-of-interest (ROI) based on the voxel-based meta-analysis by Binder and colleagues (49) (Figure 1D; heretofore referred to as “semantic network ROI”). The semantic network ROI spanned a large swathe of heteromodal cortex, including most of the aforementioned cortical areas. The neural similarity structure of concept representations in this ROI reflects not only information encoded in the activation patterns within each cortical area, but also information encoded in the pattern of activations across different areas, thus allowing us to examine semantic representation in the network as a whole.

From the fMRI data, we generated a whole-brain activation map for each concept, reflecting the unique spatial pattern of neural activity for that concept (Figure 1). From these maps, a neural representational dissimilarity matrix (RDM) was generated for each ROI. RDMs consisted of all pairwise dissimilarities (1 – correlation) between the vectorized activation patterns across voxels. For each representational model investigated, a model-based RDM was computed, and its similarity to the neural RDM was evaluated via Spearman correlation. We evaluated six different representational models: two based on taxonomic information, two based on experiential information, and two based on distributional information (see details below). The resulting profile of relative model performances provided an assessment of the degree to which each type of information is reflected in the neural activation patterns corresponding to different concepts. Because the RDMs from different models were partly correlated with each other (Table S1), we also conducted partial correlation RSAs to evaluate the unique contribution of each model to neural concept representation. We stress that the representational models investigated here are models of *information content* (i.e., taxonomic, experiential, or distributional); they are not meant to model the neural architecture through which information is encoded in the brain (see Supplementary Materials for extended model descriptions).

### Taxonomic models

*WordNet* is the most influential representational model based on taxonomic information, having been used in several neuroimaging studies to successfully model semantic content (2, 22, 54, 55). It is organized as a knowledge graph in which words are grouped into sets of synonyms (“synsets”), each expressing a distinct concept. Synsets are richly interconnected according to taxonomic relations, resulting in a hierarchically structured network encompassing 81,426 noun concepts (in the English version). Concept similarity is computed based on the shortest path connecting the two concepts in this network.

In contrast to WordNet, which provides a comprehensive taxonomy of lexical concepts, the *Categorical* model was customized to encode the particular taxonomic structure of the concept set in each study, based on a set of *a priori* categories. Therefore, the Categorical model ignores concept categories that were not included in the study, such as “furniture” or “clothing”. To reduce the level of subjectivity involved in the assignment of items to categories and in the evaluation of inter-category similarities, we tested 18 *a priori* versions of the Categorical model, with different numbers of categories and different levels of hierarchical structure, for the concepts in Study 1 (Figure S1). Each version was tested via RSA against the fMRI data, and the best performing version was selected for comparisons against other types of models. The selected model for Study 1 (model N in Figure S1) consisted of 19 hierarchically structured categories: Abstract (Mental Abstract, Social Abstract, Social Event, Other Abstract), Event (Social Event, Concrete Event), Animate (Animal, Human, Body Part), Inanimate (Artifact [Musical Instrument, Vehicle, Other Artifact], Food, Other Inanimate), and Place.

In Study 2, the Categorical model consisted of 2 higher-level categories – Object and Event – each consisting of 4 sub-categories (Animal, Plant/Food, Tool, Vehicle; Sound Event, Social Event, Communication Event, Negative Event).

### Experiential models

The *Exp48* model consists of 48 dimensions corresponding to distinct aspects of phenomenal experience, such as color, shape, manipulability, pleasantness, etc. (Table S2). This model is based on the experiential salience norms of Binder and colleagues (40), in which each dimension encodes the relative importance of an experiential attribute according to crowd-sourced ratings. Exp48 encompasses all perceptual, motor, spatial, temporal, causal, valuation, and valence dimensions present in those norms.

The *SM8* model consists of a subset of the Exp48 dimensions, focusing exclusively on sensory-motor information. These dimensions represent the relevance of each sensory modality (vision, audition, touch, taste, and smell) and of action schemas performed with each motor effector (hand, foot, and mouth) to the concept. The concept “apple”, for instance, has high values for vision, touch, taste, mouth actions, and hand actions, and low values for the other dimensions.

### Distributional models

Distributional information was modeled with two of the most prominent distributional models available: *word2vec* (56) uses a deep neural network trained to predict a word based on a context window of a few words preceding and following the target word. In contrast, *GloVe* (57) is based on the *ratio* of co-occurrence probabilities between pairs of words across the entire corpus. In a comparative evaluation of distributional semantic models (17), word2vec and GloVe emerged as the two top performing models in predicting human behavior across a variety of semantic tasks.

#### Study 1

Study 1 evaluated these six models with a set of 300 concepts spanning a wide variety of semantic categories, including animate objects (e.g., “elephant”, “student”), places (e.g., “kitchen”, “beach”), artifacts (e.g., “comb”, “rocket”) and other inanimate objects (e.g., “cloud”, “ice”), events (e.g., “hurricane”, “election”), and highly abstract concepts (e.g., “fate”, “hygiene”) (Figure 2A, Tables S3-S4).

**Figure 2:**
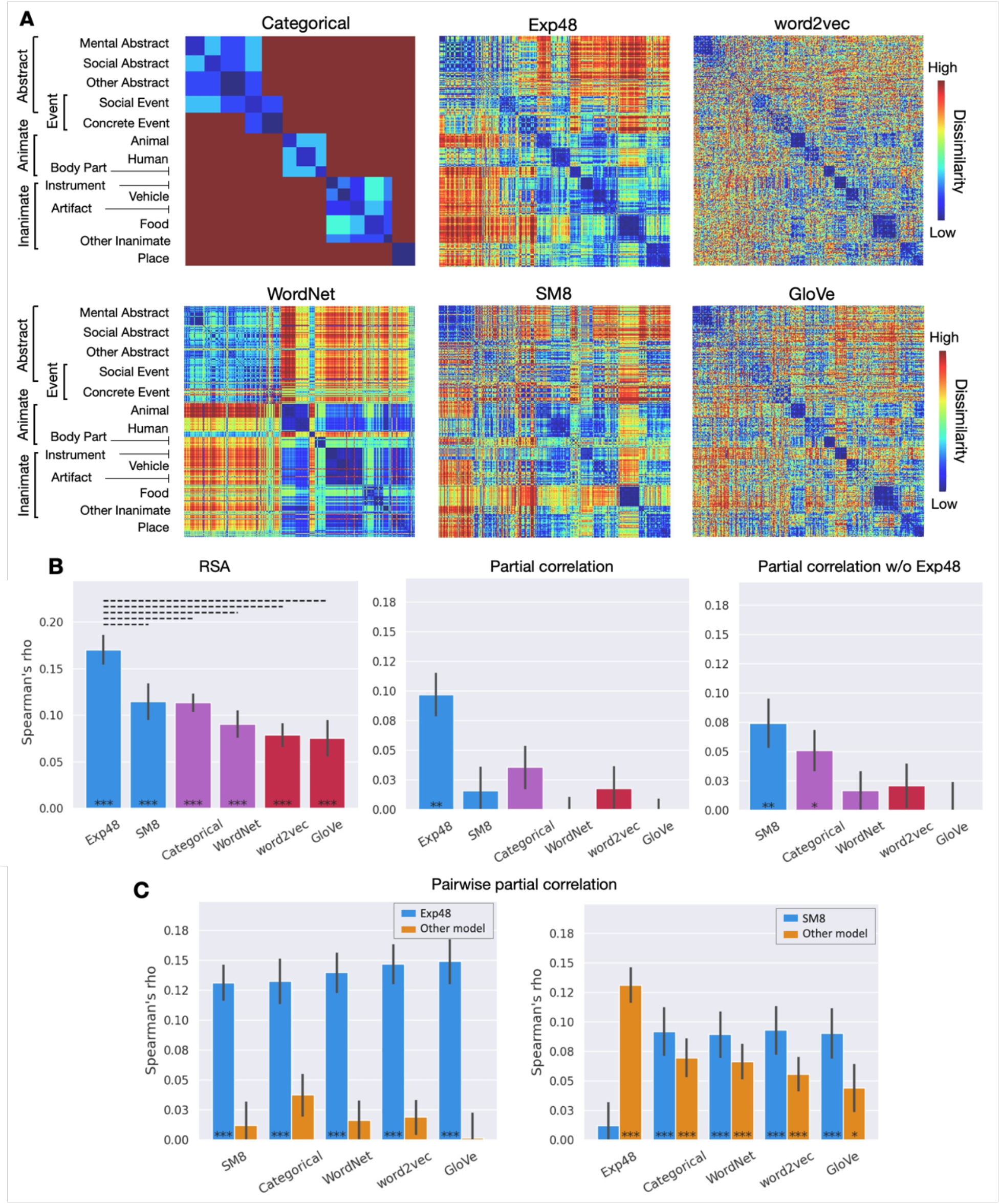
Study 1. A. Dissimilarity matrices for the representational spaces tested. Rows and columns represent each of the 300 concepts used in the study, grouped according to the Categorical model to reveal taxonomic structure. B. RSA results for the semantic network ROI. Experiential (blue), taxonomic (purple), and distributional (red) models; Left: Correlations between the group-averaged neural RDM and each model-based RDM. Center: Partial correlation results for each model while controlling for its similarity with all other models. Right: Partial correlation results when Exp48 was excluded from the analysis. C. Pairwise partial correlations for the semantic network ROI, with blue bars representing Exp48 (left) or SM8 (right) while controlling for its similarity to each of the other model-based RDMs; yellow bars correspond to each of the other model-based RDMs while controlling for their similarity to the model represented in blue. *** p < .0005, ** p < .005, * p < .05, Mantel test; solid bar: p < .001; dashed bar p < .05, permutation test. All p values are FDR-corrected for multiple comparisons (q = .05). Error bars represent the standard error.

In the functionally defined “semantic network” ROI, the experiential models achieved the highest performance, followed by the taxonomic models (Figure 2B and 2C). Exp48 performed significantly better than any other model, with no other differences between models reaching significance. To investigate the unique contribution of each type of information to the similarity structure of activation patterns, we conducted partial correlation RSAs. These analyses assessed how well each model explained the neural data after controlling for its similarity to other models. We conducted RSAs with the RDM of each model after regressing out the RDMs of all other models. As shown in Figure 2B (center), only Exp48 explained significant variance in the neural data after controlling for the other models. To verify whether Exp48 individually explains all of the variance accounted for by each of the other models, we performed pairwise partial correlation RSAs (Figure 2C, left). These analyses showed that no other model accounted for any measurable additional variance after controlling for their similarity to Exp48. In other words, Exp48 was the only model that made unique contributions to explaining the representational structure of lexical concepts in the semantic network ROI, while the other models only predicted the data to the extent that their similarity structure correlated with that of Exp48.

We then investigated whether information about the relative importance of eight sensory-motor modalities, by itself, successfully predicted the neural similarity structure of concepts after controlling for taxonomic and distributional information. The results showed that, when Exp48 was not included in the partial correlation analysis, SM8 remained highly significantly correlated with the data after controlling for all other models, while WordNet, word2vec and GloVe did not (Figure 2B, right). Categorical was the only other model displaying a significant partial correlation, although numerically lower and less significant than that of SM8. Pairwise partial correlation RSAs showed that SM8 did not explain all the variance accounted for by WordNet, word2vec, or Glove; however, there was a trend toward stronger partial correlations for SM8 than for any of the non-experiential models (Figure 2C, right). Together, these results suggest that experiential information – including but not restricted to sensory-motor information – plays a fundamental role in the neural representation of conceptual knowledge in heteromodal cortical areas, while taxonomic and distributional representational systems appear to contribute relatively little independent information.

All six models predicted the similarity structure of concept-related neural activation patterns in left AG, PCun, SFG, and IFG (Figures S2 and S3). In all of these areas, as well as in left and right PCG, left PHG, right IFG, and right SFG, the Exp48 model achieved the strongest correlation. SM8 did not perform as well as Exp48, and this difference reached significance in several ROIs, particularly in the left IFG. The same areas also showed a significant advantage for Exp48 over WordNet. The advantage of Exp48 over the Categorical model reached significance in left AG, PCun, PC, and PHG, and in right IFG and SFG. There was a trend toward higher correlations for Categorical than for WordNet in most areas, but this difference only reached significance in the left pSTS, the only area in which Categorical (or any other model) significantly outperformed both experiential models. Besides the pSTS, the Categorical model also achieved the highest correlation of any model in the left pFusG, pMTG, pSTG, and SMG, although this advantage only reached significance relative to word2vec in pSTG and pMTG and relative to SM8 in pMTG. Several areas showed a trend toward lower performance for word2vec and GloVe relative to the other models, and in most areas these two models performed similarly, with two exceptions: in left IFG, GloVe significantly outperformed word2vec, while the inverse effect was found in right PCun. The relatively low correlations for all models found in the aITG, aFusG, TP, EC, aPHG, and mOFC may be due to lower signal-to-noise ratio in those areas (Table S7).

#### Study 2

Study 2 was designed to further investigate the information content of concept representations with a different set of concepts, a larger participant sample size, and matched subsets of object and event concepts. This study examined whether the pattern of model performances found in Study 1 would be observed independently for objects and events, or – given their markedly distinct ontological status – whether these two types of concepts would differ in the degree to which they encode experiential, taxonomic, and distributional information. The 320 concepts included in the study were selected to be typical exemplars of four categories of objects (animals, tools, plants/foods, and vehicles) and four categories of events (sounds, negative events, social events, and communication events). The larger sample size (36 participants) allowed for statistical testing across participants as well as across stimuli (i.e., words).

In the semantic network ROI, Exp48 outperformed all other models when tested across participants and across stimuli (Figures 3B, left and S6, left), and, in both analyses, it was the only model that retained explanatory power after controlling for the predictions of all other models (Figure 3B, center, and S6, center). SM8 also performed significantly better than all non-experiential models, and Categorical outperformed WordNet, word2vec, and GloVe.

**Figure 3.**
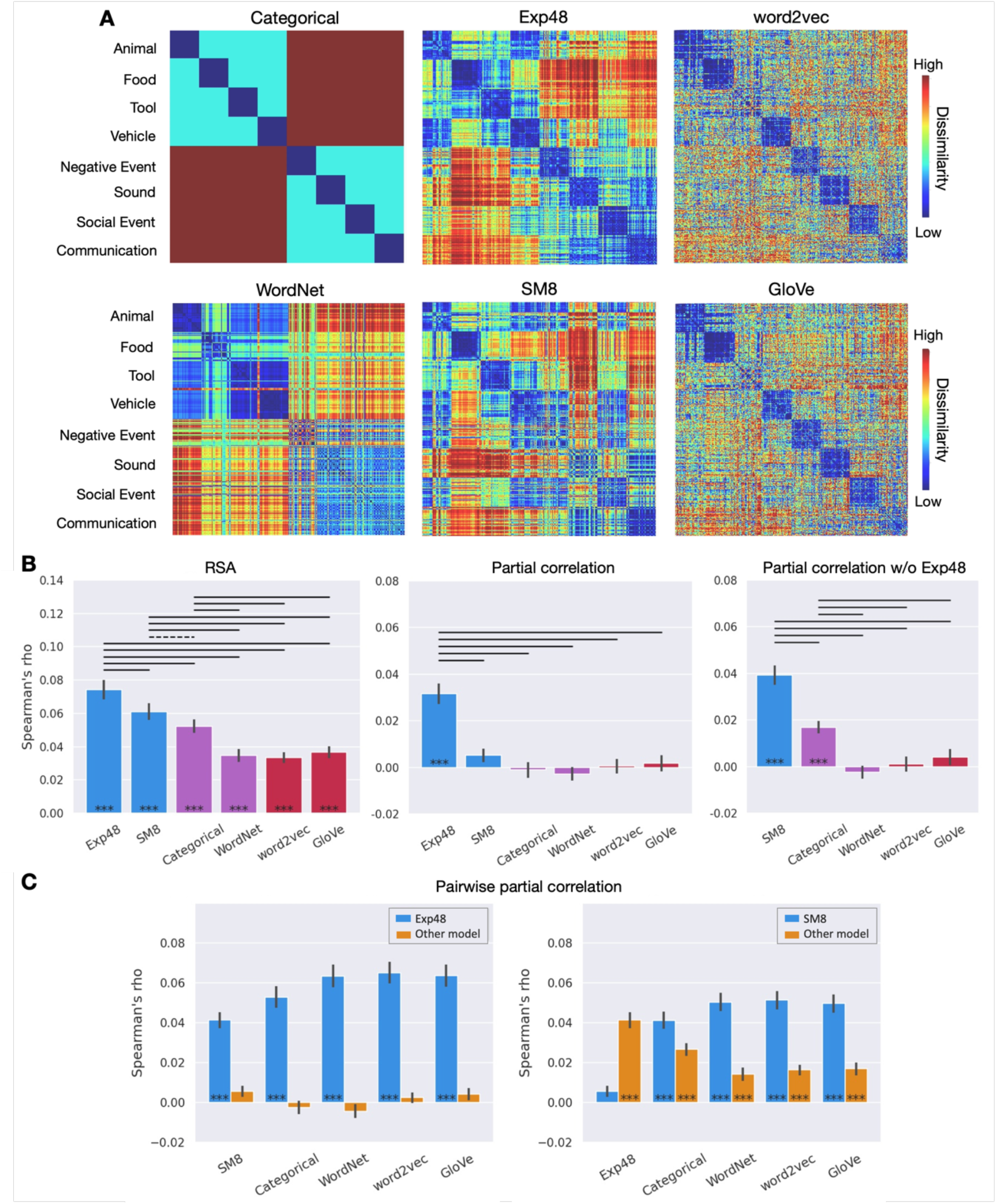
Study 2. A. Dissimilarity matrices for the representational spaces tested. Rows and columns have been grouped according to the Categorical model to reveal taxonomic structure. B. RSA results across participants for the semantic network ROI. Group mean values of the correlations between each participant’s neural RDM and the model-based RDMs. Full RSA correlations (left) and partial correlations with Exp48 included (center) and excluded (right). C. Pairwise partial correlations, with blue bars representing Exp48 (left) or SM8 (right) while controlling for its similarity to each of the other models; yellow bars correspond to each of the other models while controlling for their similarity to the model represented in blue. *** p < .0005, ** p < .005, * p < .05; solid bar: p < .001; dashed bar p < .05; Wilcoxon signed-rank tests. All p values are FDR-corrected for multiple comparisons (q = .05). Error bars represent the standard error of the mean.

Partial correlation RSAs revealed that Exp48 accounted for all the variance explained by the other models (Figure 3B, center and 3C, left). When Exp48 was left out of the analysis, SM8 and Categorical together accounted for all the variance explained (Figure 3B, right), with SM8 performing significantly better than Categorical. Confirming the main findings of Study 1, these results indicate that experiential information plays a central role in the representation of lexical concepts in high-level heteromodal cortical areas.

Separate analyses for objects and events revealed two main differences between these two types of concepts (Figures 4, S7-S10). First, RSA correlations were substantially higher for events than for objects, across all three types of representational models, in almost all ROIs. This result reflects the higher variability of pairwise similarities for the neural representations of event concepts, as evidenced by the higher noise ceiling in this condition (mean noise ceiling across anatomical ROIs, lower bound = 0.17) relative to object concepts (0.14). Inspection of the neural RDM (Figure S11) reveals a slightly more pronounced categorical structure for events than for objects, with particularly high pairwise similarities for communication events and social events.

**Figure 4.**
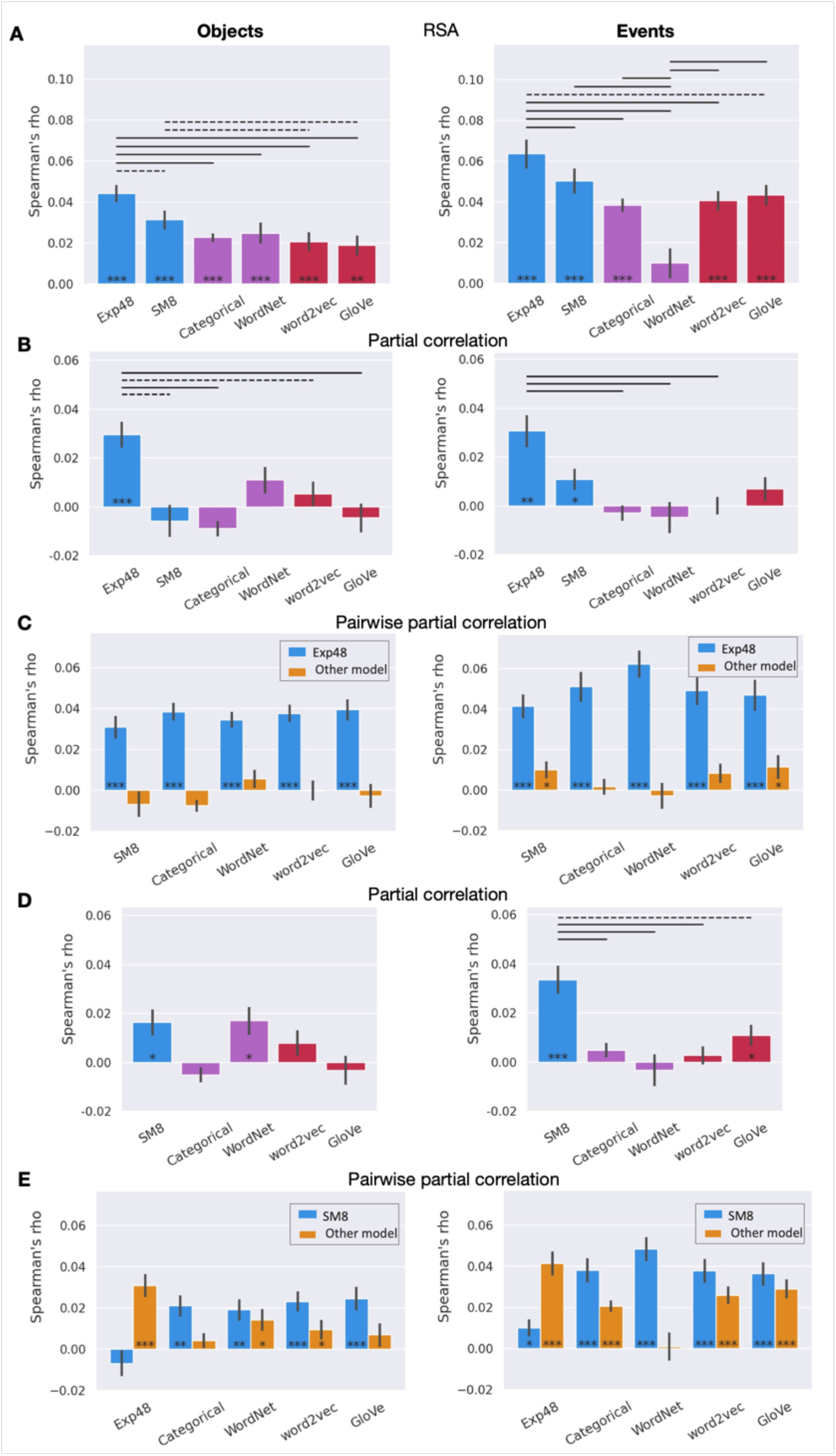
Results for object concepts (left column) and event concepts (right column) for the semantic network ROI in Study 2 (across participants). A. RSA results. B. Partial correlations. C. Pairwise partial correlations for Exp48. D. Partial correlations with Exp48 excluded from the analysis. E. Pairwise partial correlations for SM8. Color and symbol conventions as in Figure 3.

In all anatomically defined ROIs, RSAs based on the entire concept set (i.e., all categories included) revealed an advantage for experiential models relative to taxonomic and distributional models (Figures S4 and S5). As in Study 1, correlations were particularly strong in the left IFG, AG, and PCun, with the IFG showing substantially stronger correlations than other ROIs. For all models, correlations were stronger in left hemisphere ROIs than in their right hemisphere homologs, consistent with left lateralization of lexical semantic representations. The relatively low correlations found in the aITG, aFusG, TP, EC, aPHG, and mOFC may be due to lower signal-to-noise ratio in those areas (Table S7). In almost all ROIs examined (45 out of 46), correlations were significantly stronger for the experiential models than for taxonomic or distributional models. Both experiential models significantly outperformed all other models in 17 ROIs, and nowhere did a taxonomic or distributional model perform significantly better than SM8 or Exp48 (the superiority of the Categorical model in the pSTS observed in Study 1 was not replicated here). Exp48 significantly outperformed SM8 in left aSTG, pSTG, AG, pMTG, IFG, pFusG, cMFG, and SFG; and in right AG, SMG, pSTS, cMFG, and SFG. No significant differences between the two experiential models were found in the other ROIs.

Except for the left AG, WordNet appears to perform worse for events than for objects in all other ROIs, both in absolute terms and in relation to other models. This difference, which was strongest in the PCG, is likely due to taxonomic trees for events being relatively shallower in this model (mean tree depth for objects and events is 10.6 and 8.1, respectively), resulting in a narrower range of possible pairwise similarities. Apart from WordNet, the overall profile of relative model performances was similar across objects and events. For both types of concepts, experiential models achieved the strongest RSA correlations in most anatomical regions, with no ROIs showing significantly stronger correlations for any taxonomic or distributional model. Partial correlation RSAs in the semantic network ROI showed that, for object concepts, Exp48 accounted for all the variance explained by any other model (Figure 4, B and C). For events, SM8 and GloVe maintained relatively weak but statistically significant correlations after controlling for Exp48. SM8 was second to Exp48 in explaining the most variance for both objects and events (Figure 4, D and E).

#### Validation analyses

While the experiential feature ratings included in Exp48 and SM8 were meant to capture particular qualities of phenomenal experience, it is possible that the ratings themselves were influenced by other types of information as well, including more abstract knowledge about the target concept or about contextually associated concepts. Therefore, it is conceivable that a model based on semantic features that do not focus on experiential information could also outperform the taxonomic and distributional models. To assess this possibility, we conducted a new set of analyses in which we included a model based on semantic features that are not explicitly focused on experiential information. This model was based on the Semantic Feature Production Norms (SFPN), which are the largest and best-known effort to characterize word meanings in terms of descriptive semantic features (58–60). These features are derived from descriptive properties generated by human participants in a property listing task, resulting in concept representations based on thousands of features. They encode various types of information, including taxonomic (e.g., “is a mammal”), functional (e.g., “used for cooking”), and contextual association, as well as more elaborate features (e.g., “lays eggs”, “lives in the water”).

These analyses were based on the concepts included in our studies for which SFPN representations are available (220 concepts in Study 1 and 196 concepts in Study 2). The results show that SFPN was the worst performing of the seven models tested, and that both Exp48 and SM8 significantly outperformed it in both studies (Figure S12). These results indicate that the high performance of Exp48 and SM8 relative to the other models is indeed due to their focus on experiential information.

## Discussion

In two studies, we conducted a quantitative assessment of the degree to which the three main information structures proposed for concept representation are encoded in fMRI activation patterns corresponding to individual lexical concepts. We found that Exp48 – a representational model based on 48 distinct experiential dimensions grounded in known neurocognitive systems – consistently outperformed taxonomic and distributional models in predicting the neural similarity structure of a large number of lexical concepts (524 across both studies), spanning a wide variety of semantic categories. Additionally, SM8, a relatively impoverished experiential model based on only eight sensory-motor dimensions reflecting the relative importance of neural systems dedicated to vision, hearing, touch, taste, and smell, as well as motor control of the hands, mouth, and feet, exceeded the performance of the best performing taxonomic and distributional models in the semantic network ROI in Study 2. SM8 significantly outperformed those models in 17 anatomical ROIs, while matching their performance in the remaining ones. These results provide compelling evidence that experiential information grounded in sensory, motor, spatial, temporal, affective, and reward neural systems plays a fundamental role in the neural representation of concepts – not only in sensory-motor cortical areas, as indicated in previous studies, but also in high level association areas.

In both experiments, Exp48 was the only model whose predictions correlated with the neural similarity structure of the entire concept set when controlling for the predictions of all other models, and in Study 2 this was also true for object concepts in particular. For event concepts, SM8 was the only other model that independently accounted for variance in the neural data. The fact that taxonomic and distributional models did not predict the neural RDM when controlling for the predictions of Exp48 alone (Figures 2C and 3C, left panel) indicates that those models provide no measurable additional information about concept similarity structure. This suggests that taxonomic information, while seemingly central to the organization of conceptual knowledge, may be represented only indirectly, via its interdependency with experiential information (40).

Another important finding was that RSA correlations in the PCG and in the PCun were among the strongest across all ROIs examined, rivaling or even surpassing those found in the AG and in the temporal lobe. PCG and PCun are not typically included in discussions of cortical “semantic hubs” (3, 4, 28), although they have been consistently associated with semantic processing in functional neuroimaging studies (30, 49, 61, 62). This finding, replicated across Studies 1 and 2, indicates that these regions play an important and yet unrecognized role in concept representation.

One limitation of the present study is that the representational spaces used to model experiential information content are relatively coarse. Neural concept representations must encode detailed information about aspects of the phenomenal experience associated with different lexical concepts, such as particular colors, shapes, or motor schemas, but Exp48 and SM8 only encode the relative importance of each kind of experience. In other words, these models represent information about how much the neural system underlying a particular process (e.g., color perception, motor control of the hand) contributes to a lexical concept representation, but it contains no information about the particular representations contributed by each system (e.g., different hues or different motor schemas). This coarseness, which results from practical limitations in obtaining participant ratings about fine-grained experiential attributes, imposes limitations on our ability to detect experiential information in the neural data. Nevertheless, it is still surprising that these models performed so well compared to some of the most sophisticated taxonomic and distributional models available, pointing to the development of more detailed experiential models as a promising direction for future work.

Although most researchers agree that the representation of conceptual information per se should be independent of the modality of the stimulus, concept-related neural activation patterns elicited by non-verbal stimuli may differ to some extent from those elicited by words. We used words as stimuli primarily for methodological reasons: First, unlike pictures, word forms are arbitrarily associated with conceptual content. The visual, motor, or auditory representations of a word form carry no information about the semantic properties of the lexical concept associated with it. Pictorial stimuli, on the other hand, carry information about visual properties of the concept, which can bias participants to focus on those properties at the expense of properties that are not explicitly conveyed by the stimulus. Second, word stimuli allow for inclusion of relatively abstract concepts (e.g., “belief”, “year”) which are not easily conveyed via other stimulus modalities. Finally, in most experiential, taxonomic, and distributional model implementations, concepts are labeled exclusively using word forms.

Task requirements can also have an influence on activation patterns. For example, asking participants to make decisions about the stimulus based on a particular semantic criterion (e.g., living vs. non-living, large vs. small, natural vs. man-made) emphasizes those aspects of meaning to the detriment of all others. Our approach was to use a task that would be as neutral as possible regarding the conceptual content of the items. We see no reason to believe that our task favors some particular aspect of meaning over another, although that possibility has not been experimentally ruled out.

Together, the present results indicate that concept representations in heteromodal cortical areas encode information about features of phenomenal experience grounded in sensory-motor, spatiotemporal, affective, and reward systems. While other studies have shown that areas involved in sensory perception and motor control are activated during concept retrieval (13, 28, 30), our results imply that information pertaining to these systems is directly encoded in high-level association areas during concept retrieval. The idea that concepts are neurally represented as amodal symbols whose representational code is independent of modality-specific systems has a long history (23, 37); however, the finding that taxonomic and distributional models performed significantly worse than both SM8 and Exp48 in Study 2 challenges that view. While our results do not rule out the notion that concepts are organized in hierarchical taxonomic networks, they are more aligned with a view in which concept representations emerge from the multimodal integration of signals originating in modality-specific systems during concept acquisition, and in which taxonomic organization emerges from correlations between experiential features (14, 30). In this framework, concept retrieval consists of the transient activation of a neuronal cell assembly distributed across the heteromodal cortical hubs that make up the semantic network, with possible downstream activation of the relevant modality-specific assemblies depending on contextual demands. Further research is required to determine the extent to which different experiential features contribute to concept representation in different parts of the semantic network, and how their relative importance varies across ontological categories.

In addition to their implications for the nature of the representations underlying conceptual knowledge, the present results are also relevant to computational approaches to natural language processing in the field of artificial intelligence. They suggest that incorporating experiential information into computational models of word meaning could lead to more human-like performance. These findings can also inform the development of brain-machine interface systems by demonstrating that information about the experiential content of conceptual thought can be decoded from the spatial distribution of neural activity throughout the cortex.

## Materials and Methods

### Study 1

This study was designed for RSA across stimuli rather than across participants, with a large number of trials (1,800 trials) and long scanning times, which allowed sufficient power to be achieved with a relatively small sample size (N = 8).

#### Participants

Eight native speakers of English (4 women), ages 19-37 (mean = 28.5) took part in Study 1. Participants were all right-handed according to the Edinburgh Handedness Scale (63), had at least a high school education, and had no history of neurological or psychiatric conditions. All participants provided written informed consent. This study was approved by the Medical College of Wisconsin Institutional Review Board.

#### Stimuli

Stimuli included 300 English nouns of various categories, including concrete and abstract concepts. Nouns were relatively familiar, with mean log-transformed HAL frequency of 8.7 (range 4.0–12.7) and mean age of acquisition of 6.7 years (range 2.7–13.8; English Lexicon Project (https://elexicon.wustl.edu) (64); Tables S3-S4).

#### Task

Participants rated each noun according to how often they encountered the corresponding entity or event in their daily lives, on a scale from 1 (“rarely or never”) to 3 (“often”). The task was designed to encourage semantic processing of the word stimuli without emphasizing any particular semantic features or dimensions. Participants indicated their response by pressing one of three buttons on a response pad with their right hand. On each trial, a noun was displayed in written form on the center of the screen for 500 ms, followed by a 3.5-second blank screen. Each trial was followed by a central fixation cross with variable duration between 1 and 4 s (mean = 2 s). The entire stimulus set was presented 6 times over the course of the study. In each presentation, the order of the stimuli was randomized. Each presentation was split into 3 runs of 100 trials each. The task was performed over the course of 3 scanning sessions on separate days, with 2 presentations (6 runs) per session. Each run started and ended with an 8-second fixation cross.

#### Equipment

Scanning was performed on a GE Healthcare Discovery MR750 3T MRI scanner at the Medical College of Wisconsin’s Center for Imaging Research. Stimulus presentation and response recording were performed via E-prime 2.0 software running on a Windows desktop computer and a Psychology Software Tools Serial Response Box. Stimuli were back-projected on a screen positioned behind the scanner bore and viewed through a mirror attached to the head coil.

#### Scanning protocol

MRI scanning was conducted over 3 separate sessions on different days. Each session consisted of a structural T1-weighted MPRAGE scan, a structural T2-weighted CUBE scan, 3 pairs of T2-weighted spin echo echo-planar scans (5 volumes each) acquired with opposing phase-encoding directions (for correction of geometrical distortions in the functional images), and 6 gradient echo echo-planar runs using simultaneous multi-slice acquisition (8x multiband, TR = 1200 ms, TE = 33.5 ms, 512 volumes, flip angle = 65, matrix = 104 × 104, slice thickness = 2.0 mm, axial acquisition, 72 slices, field-of-view = 208 mm, voxel size = 2 × 2 × 2 mm).

#### Data analysis

Functional MRI images were pre-processed (slice timing correction, motion correction, distortion correction, volume alignment, scaling) using AFNI (65). Statistical analyses were conducted in each participant’s original coordinate space. Activation (beta) maps were generated for each noun relative to the mean signal across all other nouns using AFNI’s 3dDeconvolve. Response time z scores and head motion parameters were included as regressors of no interest. Anatomically defined ROIs were created based on the probabilistic Desikan-Killiany parcellation atlas included in AFNI (TT_desai_dk_mpm; Figure S12). The functionally defined semantic network ROI corresponded to the 1% most consistently activated voxels in an activation likelihood estimate (ALE) meta-analysis of 120 neuroimaging studies of semantic language processing (49) (main text Figure 1D). All ROI masks were created in MNI coordinate space and non-linearly aligned to each participant’s anatomical scan using AFNI’s 3dQwarp. Neural RDMs were computed for each ROI as the Spearman correlation distance (1 – rho) for all pairs of concepts. A group-averaged neural RDM was computed by averaging the neural RDMs of all participants. A model-based RDM was computed from each model (except WordNet) as the cosine distance (1 – cosine) for all pairs of concept vectors. The WordNet RDM was based on the Wu & Palmer similarity metric (WPsim), which relies on the depth of the two synsets in the taxonomic tree and that of their Least Common Subsumer (LCS, i.e., their most specific common hypernym):

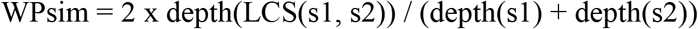

We used the package Natural Language Toolkit (NLTK 3.4.5; https://www.nltk.org) to compute WPsim; WordNet dissimilarity was computed as 1 – WPsim.

The RSAs computed the Spearman correlation between the group-averaged neural RDMs and each model-based RDM. Statistical significance was tested using the Mantel test with 10,000 permutations. Differences in RSA correlations between models were assessed for significance via permutation tests. Significance was controlled for multiple comparisons via false discovery rate (FDR) with q = .05. Since the model-based RDMs can be strongly correlated between models (Table S1), these analyses were supplemented with partial correlation analyses to reveal the unique contribution of each model after controlling for other models.

We also conducted a temporal signal-to-noise ratio (tSNR) analysis to assess data quality across the cortical mantle. A GLM was conducted with parametric regressors for word concreteness, word length, response time, and head motion parameters, and a binary regressor for trial onset. Each regressor was convolved with a canonical HRF to generate the design matrix. A GLM was conducted on the scaled, smoothed (6 mm) imaging data for each participant, and a tSNR map was computed by dividing the mean signal by the standard deviation of the residuals. The individual maps were warped into the MNI 152 2009c template via affine transformation and averaged together to generate a group-averaged tSNR map (Figure S13).

### Study 2

Study 2 was designed to assess the contribution of each representational system separately for object and event concepts in the semantic network ROI, and to achieve sufficient power for model comparisons within anatomically defined ROIs with RSA across participants as well as across stimuli, which required a substantially larger sample size (N = 36).

#### Participants

36 native speakers of English (21 women), ages 20-41 (mean = 28.7) took part in Study 2. Participants were all right-handed according to the Edinburgh Handedness Scale (63), had at least a high school education, and had no history of neurological or psychiatric conditions. None of the participants took part in Study 1. All participants provided written informed consent. This study was approved by the Medical College of Wisconsin Institutional Review Board.

#### Stimuli

Stimuli included 160 object nouns (animals, foods, tools, and vehicles – 40 of each) and 160 event nouns (social events, verbal communication events, non-verbal sound events, negative events – 40 of each) (Tables S5-S6). Of the 320 concepts included in Study 2, 62 objects and 34 events were also used in Study 1.

#### Task

Task instructions were identical to Study 1. On each trial, a noun was displayed in written form on the center of the screen for 500 ms, followed by a 2.5-second blank screen. Each trial was followed by a central fixation cross with variable duration between 1 and 3 s (mean = 1.5 s). The entire stimulus set was presented 6 times over the course of the study in randomized order. Each presentation was split into 4 runs of 80 trials each. The task was performed over the course of 3 scanning sessions on separate days, with 2 presentations (8 runs) per session. Each run started and ended with an 8-second fixation cross.

#### Equipment

Scanning was performed on a GE Healthcare Premier 3T MRI scanner with a 32-channel Nova head coil at the Medical College of Wisconsin’s Center for Imaging Research. Stimulus presentation and response recording were performed with Psychopy 3 software (66) running on a Windows desktop computer and a Celeritas fiber optic response system (Psychology Software Tools, Inc.). Stimuli were displayed on an MRI-compatible LCD screen positioned behind the scanner bore and viewed through a mirror attached to the head coil.

#### Scanning protocol

MRI scanning was conducted on 3 separate visits. Each session consisted of a structural T1-weighted MPRAGE scan, a structural T2-weighted CUBE scan, 3 pairs of T2-weighted spin echo echo-planar scans (5 volumes each) acquired with opposing phase-encoding directions, and 8 gradient echo echo-planar functional scans (4x multiband, TR = 1500 ms, TE = 33 ms, 251 volumes, flip angle = 50, in-plane matrix = 104 × 104, slice thickness = 2.0 mm, axial acquisition, 68 slices, field-of-view = 208 mm, voxel size = 2 × 2 × 2 mm).

#### Data analysis

Data preprocessing and statistical analysis were performed as described in Study 1. RDMs for the models tested in Study 2 are depicted in Figure 3 (main text), and inter-model correlations are displayed in Table S1.

RSAs were conducted across stimuli (i.e., words) and across participants. In the analysis across stimuli, voxel-based RDMs from all participants were combined into a group-averaged neural RDM, and, for each model, a single correlation was computed between this RDM and the model-based RDM. These correlations were tested for significance via the Mantel test with 10,000 permutations. In the analysis across participants, a correlation was computed, for each model, between each participant’s neural RDM and the model-based RDM. The Fisher Z-transformed correlation coefficients were then averaged across participants to compute the group mean rho, and significance tested via Wilcoxon’s Signed Rank test.

Differences in RSA correlations between models were tested across words via permutation test (Mantel) and across participants via Wilcoxon’s Signed-Rank test. Significance was controlled for multiple comparisons via false discovery rate (FDR) with q = .05.

A group-averaged tSNR map was computed as in Study 1.

## Acknowledgments

The authors wish to thank Elizabeth Awe and Jed Mathis for assistance with data collection and management and two anonymous reviewers for valuable comments and suggestions on a previous version of the article.

## Author contributions

L.F. conceived the study, coordinated data acquisition, analyzed the data, prepared figures and tables, and wrote the initial draft; J. Tong contributed to data analyses; C.J.H. contributed software tools for data acquisition; J.R.B. and L.L.C. contributed to task design, developed the stimulus set, and reviewed/edited the manuscript; J.R.B supervised the project and acquired financial support.

