## Supplemental Materials for "Decoding the Information Structure Underlying the Neural Representation of Concepts"

**This PDF file includes:**

Supplementary Text
Figs. S1 to S13
Tables S1 to S7
Supplementary References

### Supplementary Information Text

Representational Models

***WordNet*** (1) is a lexical database in which words are grouped into sets of synonyms (synsets), each expressing a distinct concept. Word forms with several distinct meanings (homonyms and polysemous words) are represented in as many distinct synsets. Synsets are interconnected according to conceptual-semantic relations. Our WordNet model is based on the superordinate-subordinate relation (hypernymy-hyponymy), which links more general synsets (e.g., “vehicle”) to increasingly specific ones (e.g., “car” and “sedan”). Thus, the WordNet model encodes hierarchical taxonomic information about lexical concepts, such as that the category “vehicle” includes “car”, which in turn includes “sedan”; conversely, concepts like “car” and “sedan” make up the category vehicle. This hierarchical structure is represented as a tree, and all noun hierarchies ultimately go up to the root node (“entity”). We used the Natural Language Toolkit (NLTK 3.4.5; https://www.nltk.org) to compute WordNet concept similarity. NLTK implements several methods for computing representational similarity between synsets. We report here the results obtained via the Wu-Palmer method, which achieved the highest RSA performance for WordNet-based RDMs in a preliminary analysis. The method is based on the depth of the two synsets in the taxonomic tree and that of their Least Common Subsumer (LCS, i.e., their most specific common hypernym).

The ***Categorical*** model encoded superordinate-subordinate relations exclusively for the concepts in each study. It was customized to the stimulus set, such that the categories were chosen to fit the particular set of concepts included in each study. Unlike WordNet, which is structured as a deep taxonomic tree connecting all noun concepts, the Categorical model represented concepts in a shallow tree, consisting of three (Study 1) or two (Study 2) levels. The model included one binary factor (yes/no) per taxonomic category. In Study 1, the model consisted of 19 hierarchically structured categories: Abstract (Mental Abstract, Social Abstract, Social Event, Other Abstract), Event (Social Event, Concrete Event), Animate (Animal, Human, Body Part), Inanimate (Artifact [Musical Instrument, Vehicle, Other Artifact], Food, Other Inanimate), and Place. Concept vector representations were, thus, 19-dimensional. In Study 2, the model consisted of 2 higher-level categories – Object and Event – each consisting of 4 sub-categories (Animal, Food, Tool, and Vehicle; Sound Event, Social Event, Communication Event, and Negative Event), resulting in 10-dimensional vector representations (Figure 2).

The ***Exp48*** model consists of 48 dimensions corresponding to distinct types of human phenomenal experience (Table S2). The model is based on the experiential salience norms by Binder and colleagues (5). Each dimension encodes the relative importance of an experiential domain according to crowd-sourced ratings on a Likert-type scale obtained via the Amazon Mechanical Turk (AMT) online platform. An important criterion for the inclusion of domains was that they could be mapped onto independently established neurocognitive processes (i.e., processes operationalized independently of semantic tasks). Seventeen components of the experiential salience norms were not included in Exp48: To avoid introducing taxonomic information into the model, we excluded features that, although related to sensory perception, are strongly associated with particular semantic categories, such as Face and Body (humans), Speech and Communication (human communication events), Music (musical instruments), and Biomotion (humans and animals). We also left out features that have not been clearly operationalized independently of conceptual knowledge, such as Cognition, Self, Complexity and Number. Finally, since emotional/affective/interoceptive components are represented in the experiential ratings both in terms of emotion categories (Happy, Sad, Angry, Disgusted, Fearful, Surprised) and of more elementary affective and reward features (Benefit, Harm, Pleasant, Unpleasant, Drive, Needs, Arousal), we included only the latter. The experiential ratings are available from https://www.neuro.mcw.edu/index.php/resources/brain-based-semantic-representations.

The ***SM8*** model consists of a subset of 5 sensory (vision, audition, touch, taste, smell) and 3 motor (hand, foot, mouth) dimensions from Exp48. These dimensions are meant to capture information about the relative level of contribution of each of the major sensory perception domains, and of somato-motor control systems for each of the three main motor effectors, to concept representations stored in multimodal/heteromodal cortex.

***word2vec*** (2) is a distributional model that, rather than directly computing word co-occurrence frequencies, uses a deep neural network trained to predict a word based on its local context. Unlike Latent Semantic Analysis (LSA), which relies on the global context in which a word occurs, word2vec relies on a context window of a few words preceding and following the target word. In a comparative evaluation of semantic word embeddings (3), word2vec emerged as one of the two top performing models (along with GloVe) in predicting human behavior across a variety of semantic tasks. We used the 300-dimensional word vectors trained on the Google News dataset (approximately 100 billion words) based on the continuous skip-gram algorithm and distributed by Google (https://code.google.com/archive/p/word2vec).

***GloVe*** (4) is a distributional model designed to learn word vectors such that their dot product equals the logarithm of the words' probability of co-occurrence. In contrast to LSA, which directly encodes word co-occurrence probabilities, and to word2vec, which learns to predict words based on their local context, GloVe is based on the *ratio* of co-occurrence probabilities between pairs of words across the entire corpus. It was shown to outperform word2vec on a word analogy task and in several word similarity tasks (4), but the two models performed similarly in the tasks analyzed by Pereira and colleagues (3). We used the 300-dimensional word vectors trained on Common Crawl (840 billion words) and made available by the authors (https://nlp.stanford.edu/projects/glove).

The ***Semantic Feature Production Norms*** **(SFPN)** devised by McRae and colleagues (12) are arguably the largest and best-known effort to characterize word meanings in terms of semantic features, and have been used to test a variety of claims about the organization of the semantic system (13–15). Features are derived from descriptive properties generated by human participants in a property listing task. The properties are subsequently standardized into features with a binary value (present/absent), and their frequencies computed for each concept. This procedure results in vector-based concept representations based on thousands of features. The features represent various types of information, including perceptual (e.g., “is red”, “roars”), taxonomic (e.g., “is a mammal”), functional (e.g., “used for cooking”), and contextual association, as well as more concept-specific types of information (e.g., “lays eggs”, “lives in the water”). We used the list of cosine similarities provided by Buchanan and colleagues (16) to generate the SFPN RDM (downloaded from https://github.com/doomlab/Word-Norms-2).

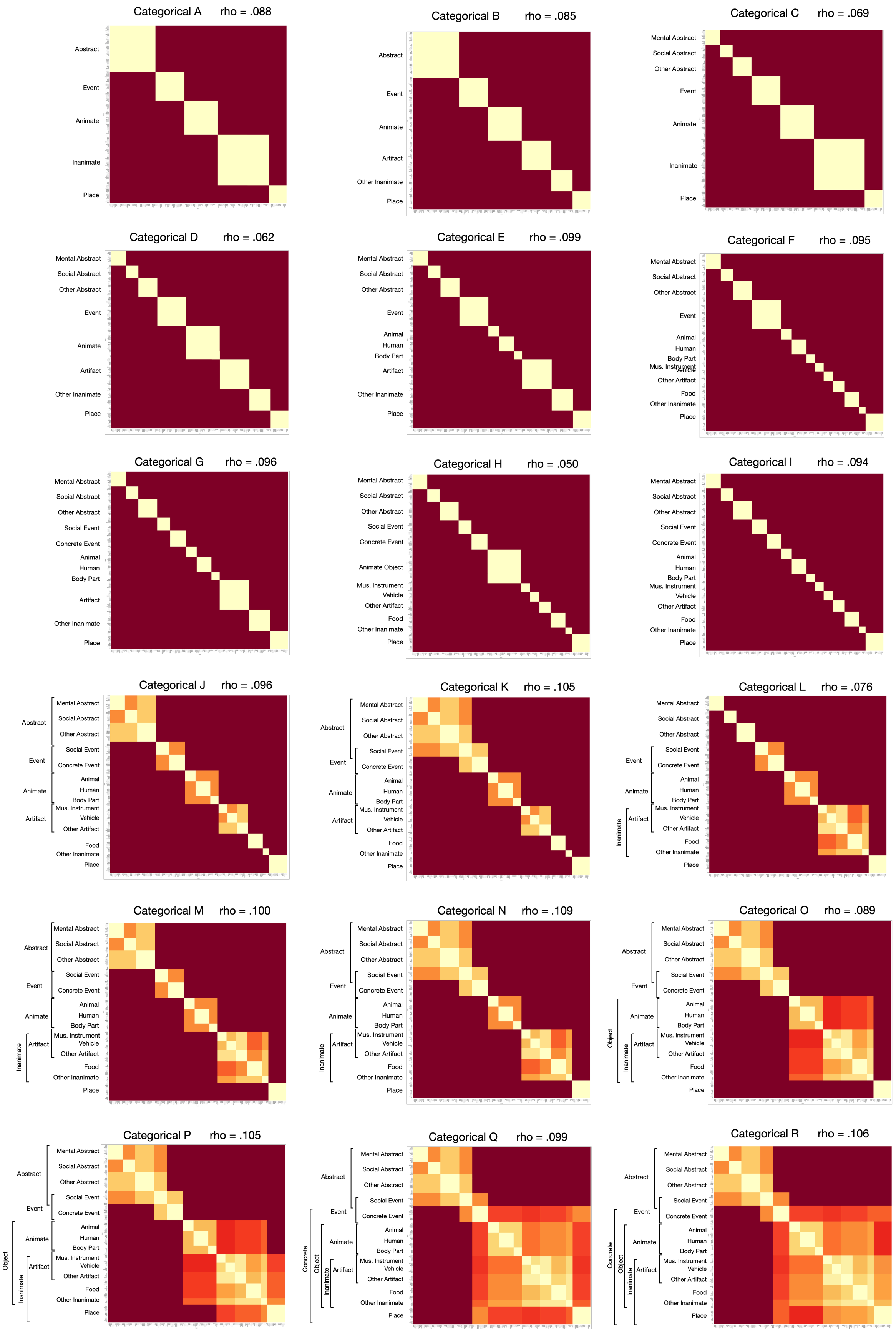
Categorical A rho = .088
Categorical B rho = .085
Categorical C rho = .069
Categorical D rho = .062
Categorical E rho = .099
Categorical F rho = .095
Categorical G rho = .096
Categorical H rho = .050
Categorical I rho = .094
Categorical J rho = .096
Categorical K rho = .105
Categorical L rho = .076
Categorical M rho = .100
Categorical N rho = .109
Categorical O rho = .089
Categorical P rho = .105
Categorical Q rho = .099
Categorical R rho = .106

**Figure S1.** Categorical models tested for Study 1. Taxonomic structure, representational similarity matrix, and Spearman's rho (computed for the semantic network ROI) for each version tested. The best performing model (N) was selected for the main analyses.

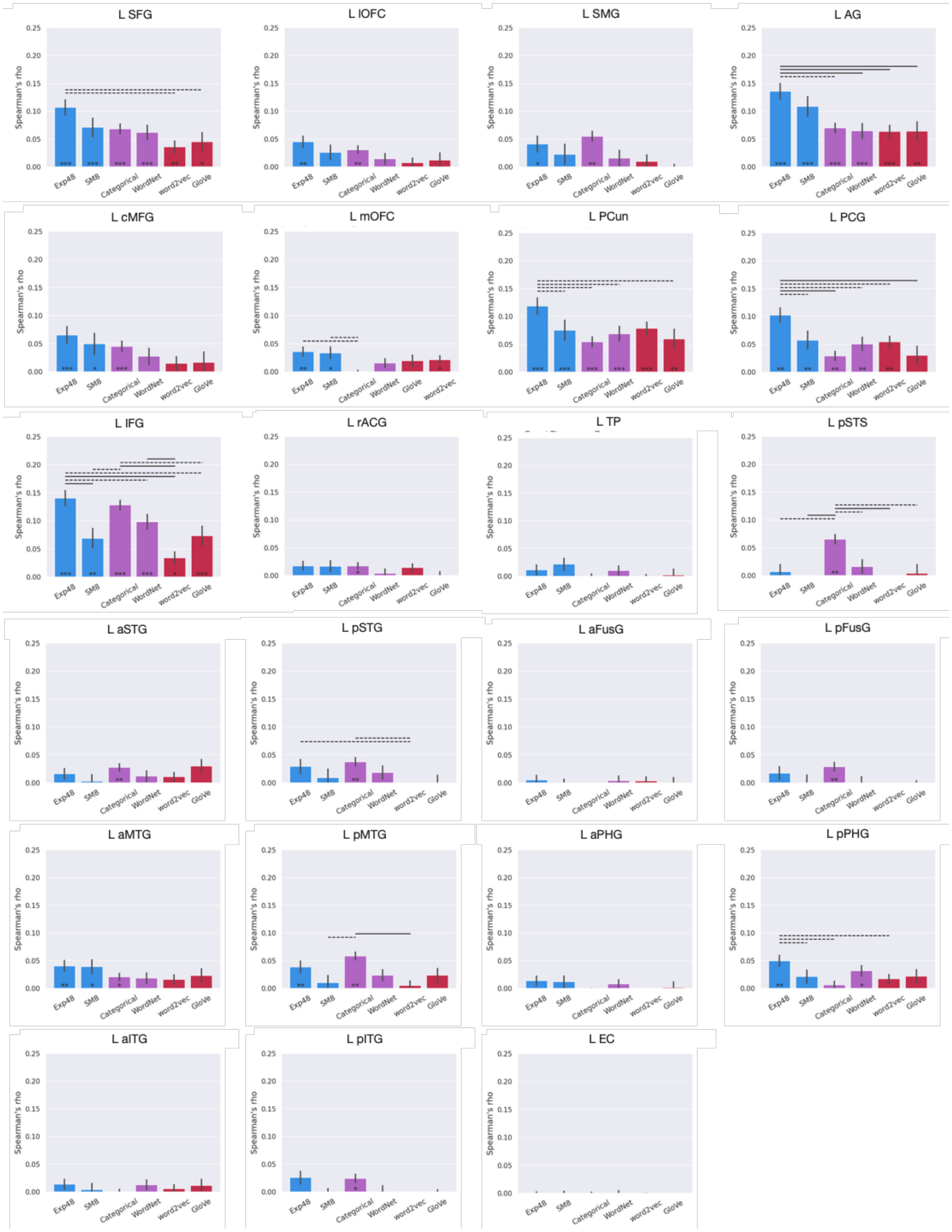


**Figure S2**. RSA results (across stimuli) for left hemisphere anatomical ROIs in Study 1. Color and symbol conventions as in Figure 2.

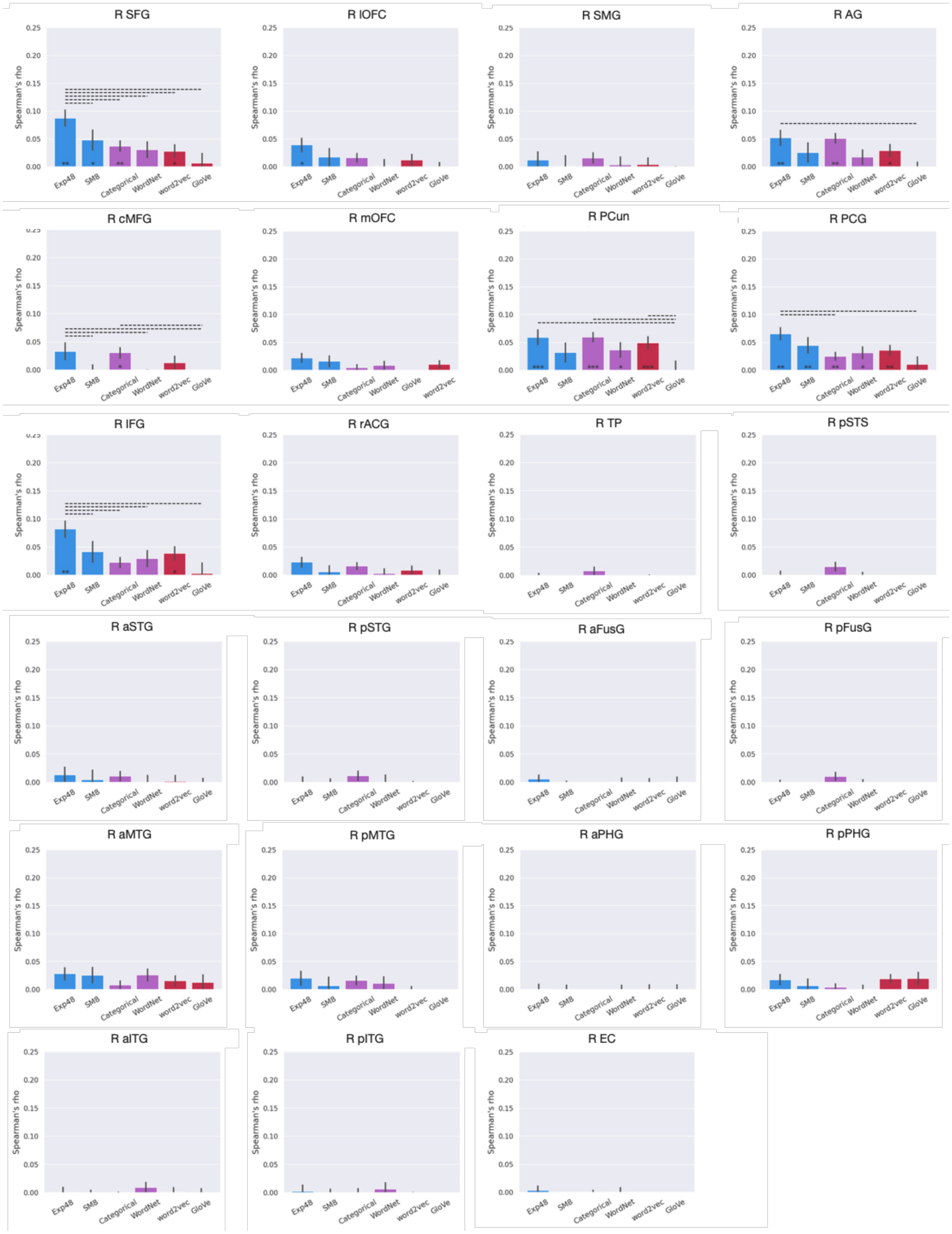


**Figure S3.** RSA results (across stimuli) for right hemisphere anatomical ROIs in Study 1. Color and symbol conventions as in Figure 2.

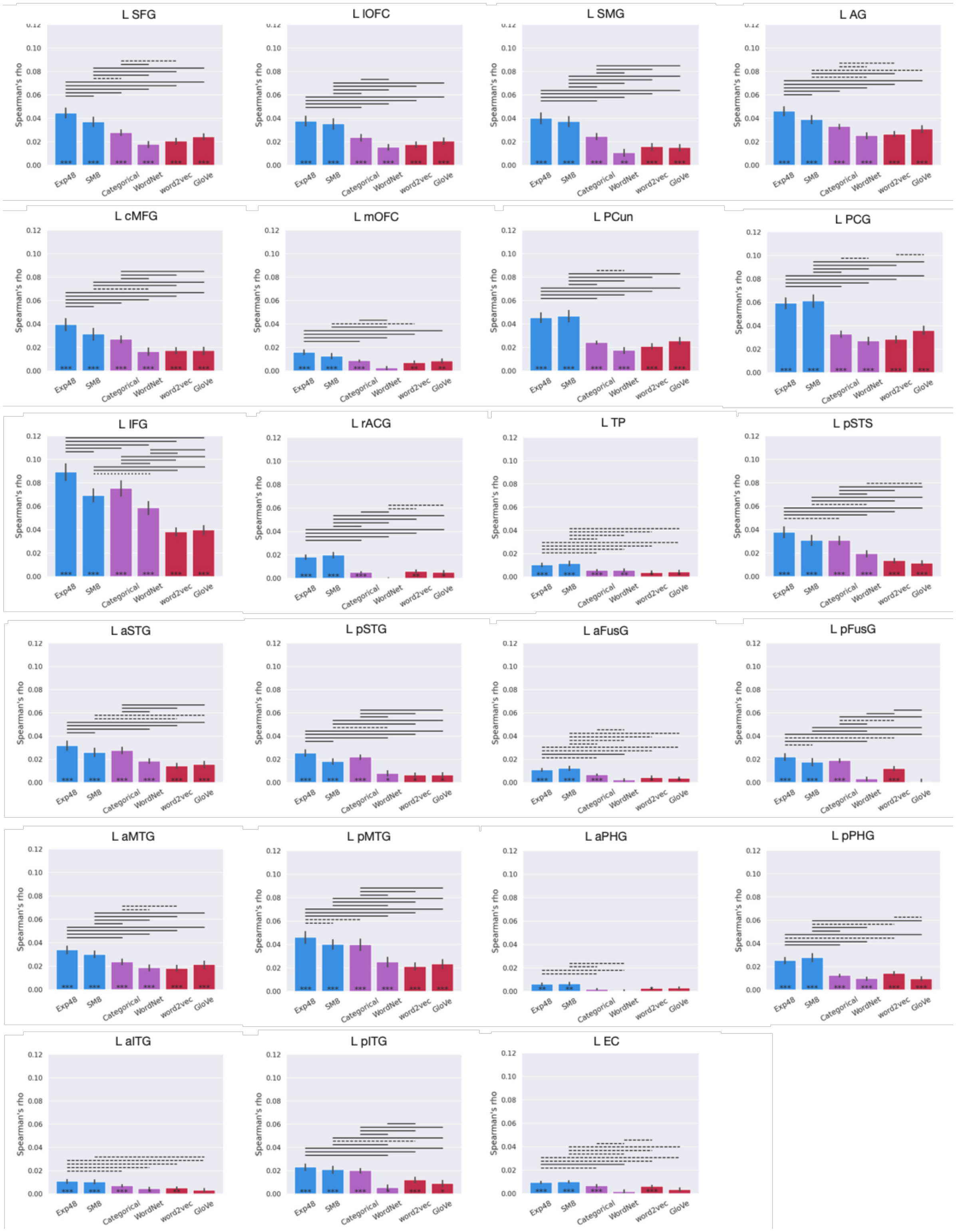


**Figure S4.** RSA results (across participants) for left hemisphere anatomical ROIs in Study 2. Color and symbol conventions as in Figure 3.

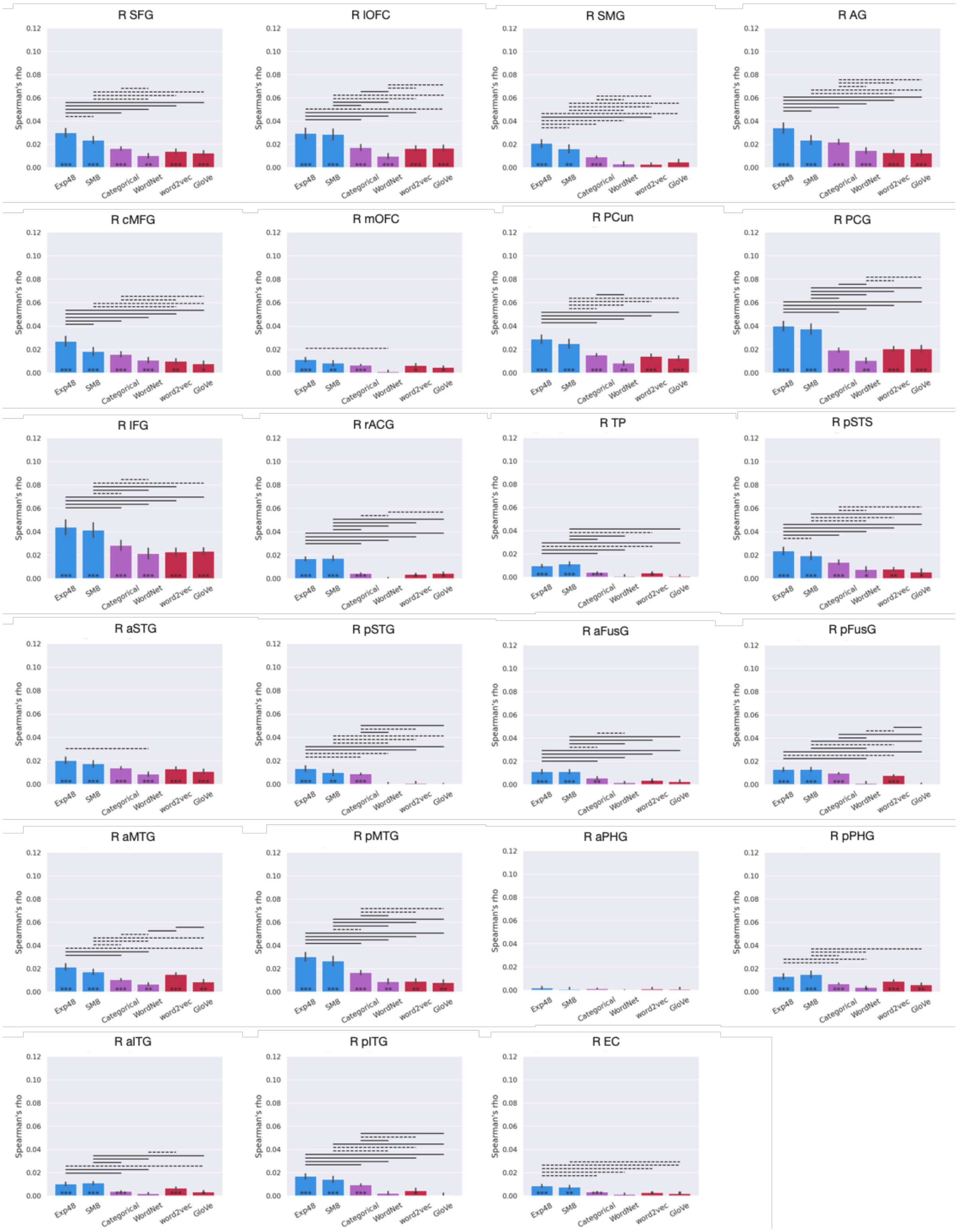


**Figure S5.** RSA results (across participants) for right hemisphere anatomical ROIs in Study 2. Color and symbol conventions as in Figure 3.

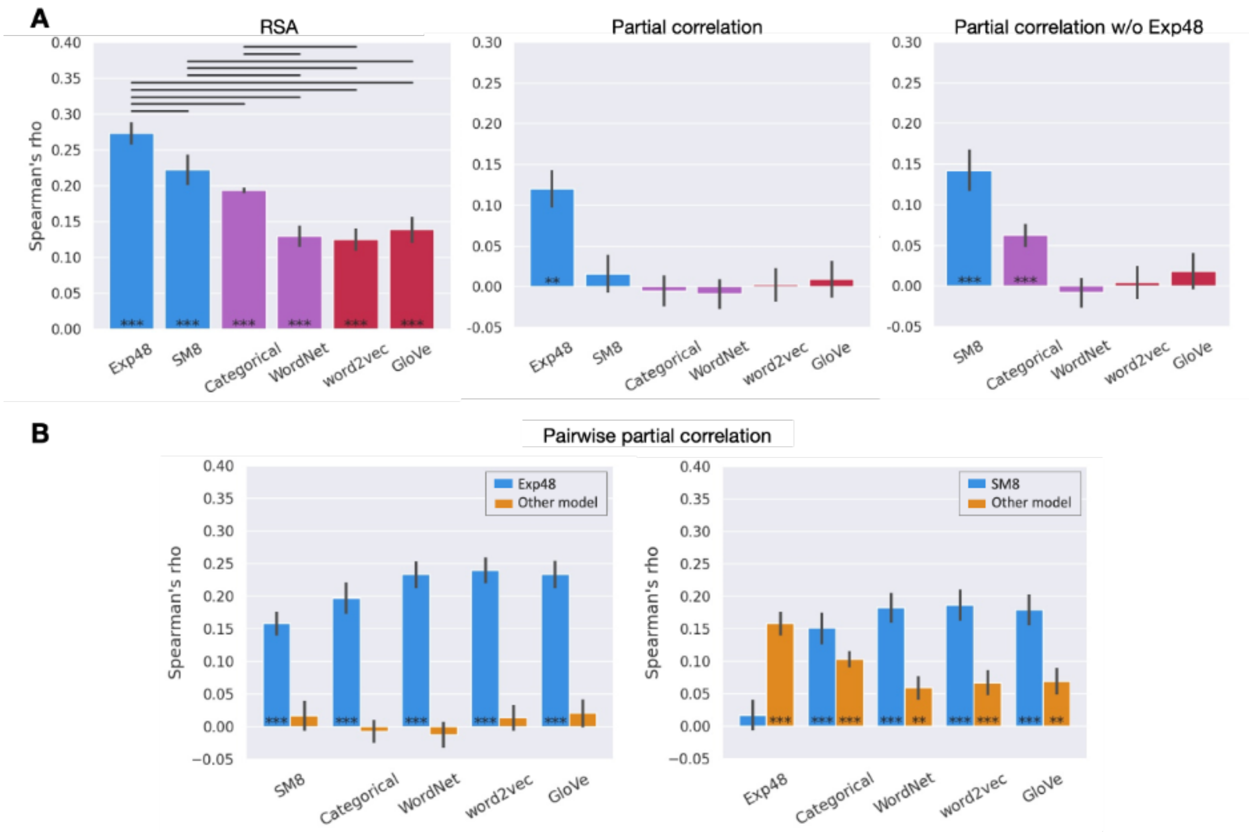


**Figure S6.** RSA results (across stimuli) for the semantic network ROI in Study 2. A. Experiential (blue), taxonomic (purple), and distributional (red) models; Left: Correlations between the group-averaged neural RDM and each model-based RDM. Center: Partial correlation results for each model while controlling for its similarity with all other models. Right: Partial correlation results when Exp48 was excluded from the analysis. B. Pairwise partial correlations for the semantic network ROI; blue bars represent Exp48 (left) or SM8 (right) while controlling for its similarity to each of the other model-based RDMs. Color and symbol conventions as in Figure 2.

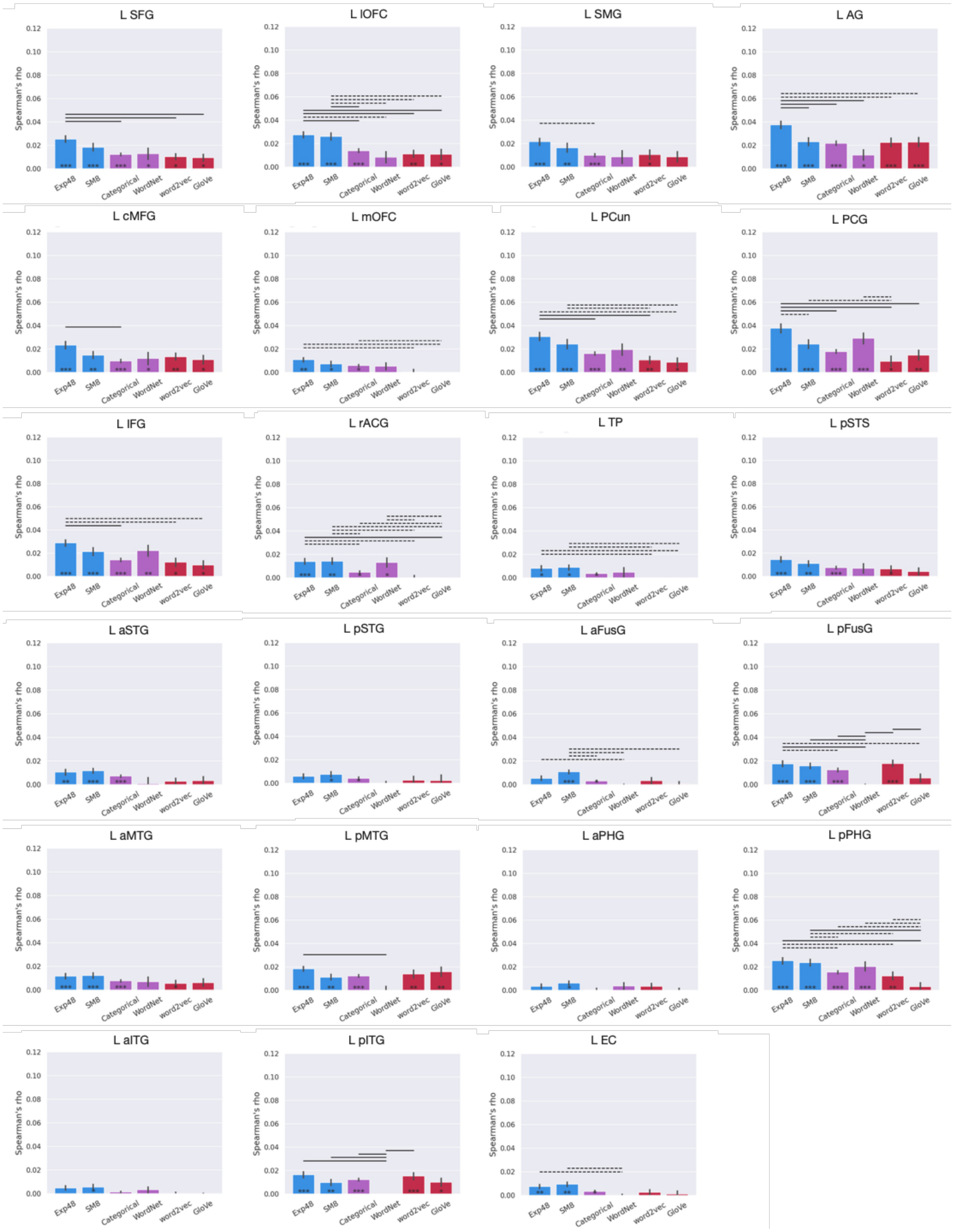


**Figure S7.** RSA results (across participants) for object concepts (left hemisphere anatomical ROIs). Color and symbol conventions as in Figure 3.

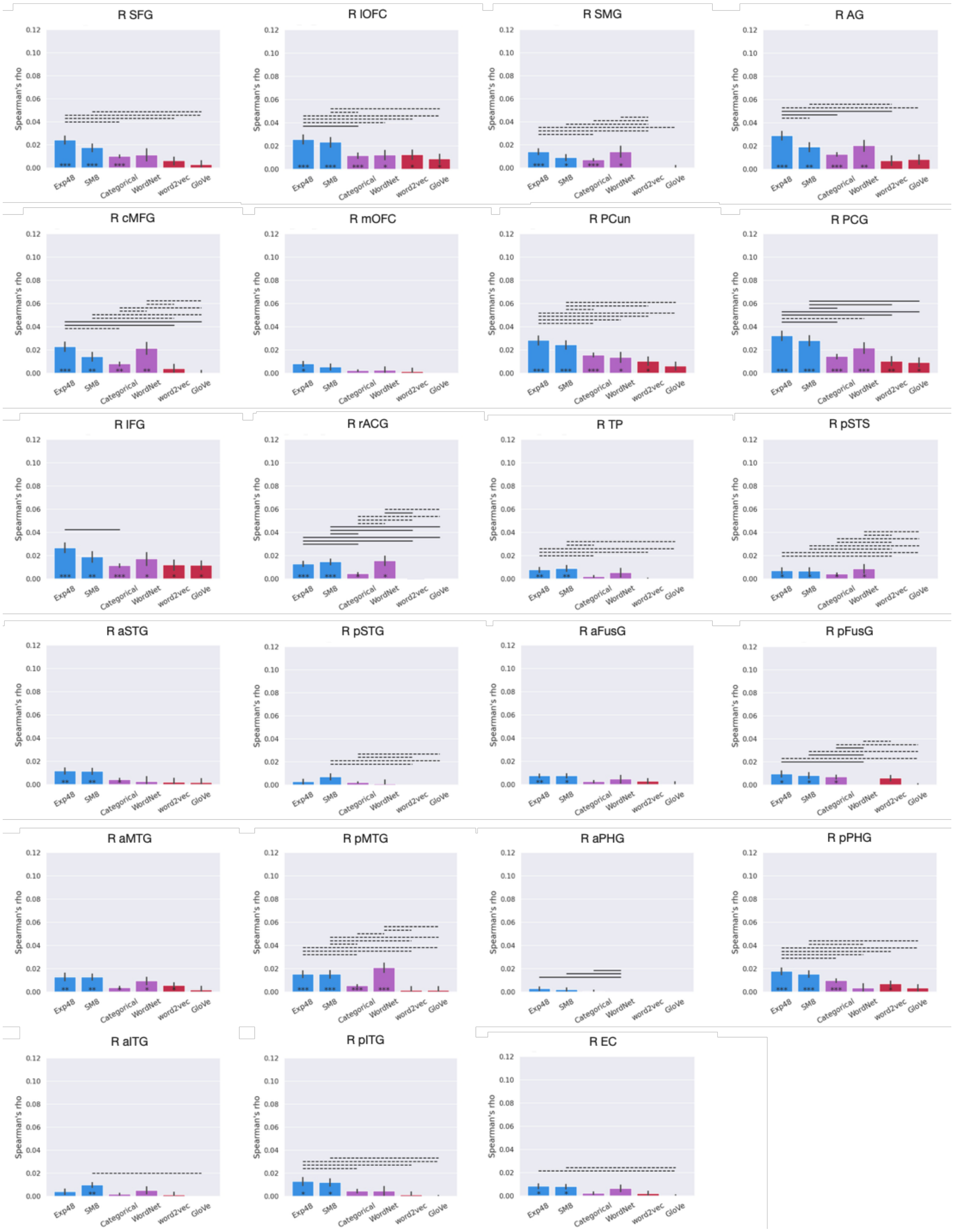


**Figure S8**. RSA results (across participants) for object concepts (right hemisphere anatomical ROIs). Color and symbol conventions as in Figure 3.

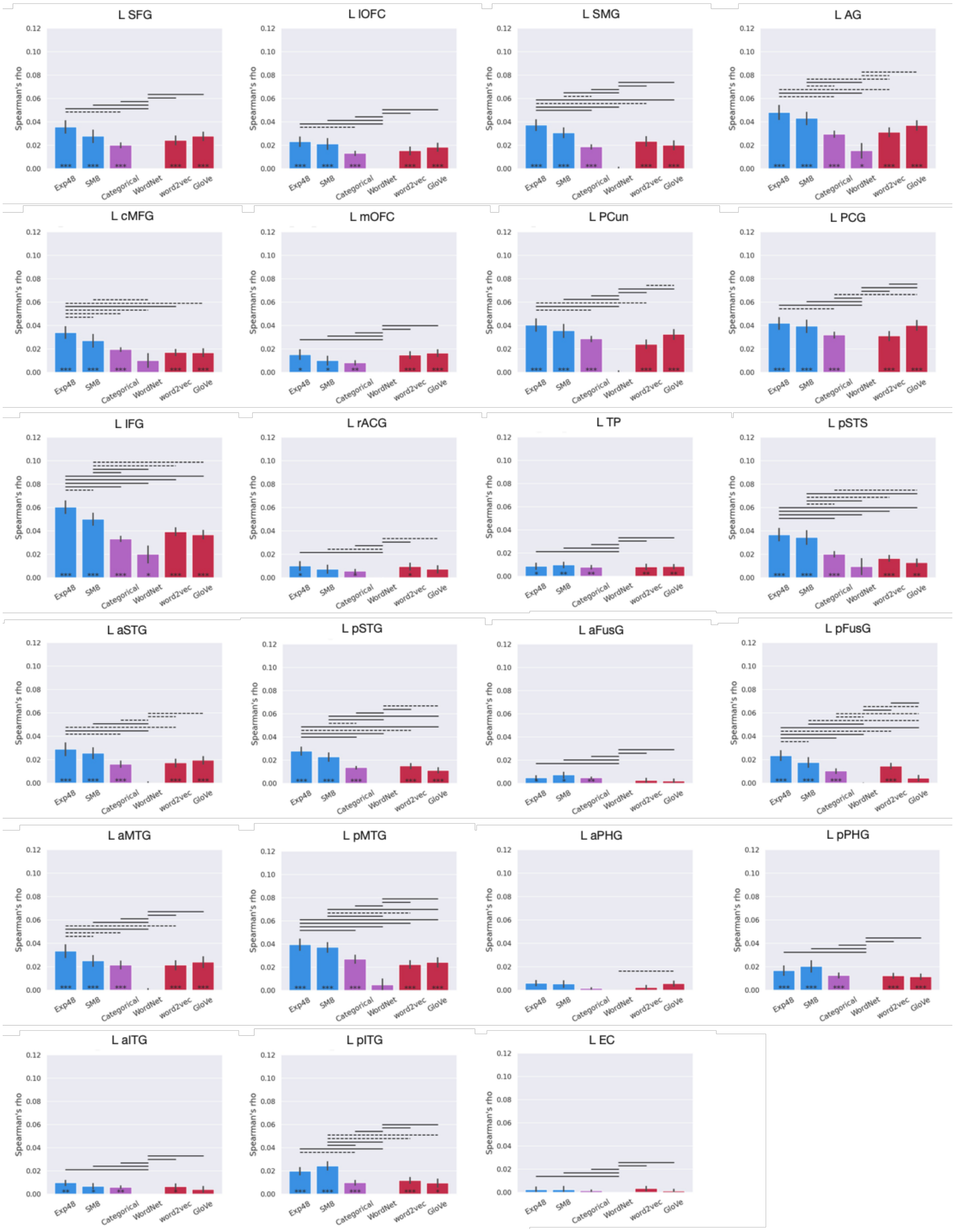


**Figure S9.** RSA results (across participants) for event concepts (left hemisphere anatomical ROIs). Color and symbol conventions as in Figure 3.

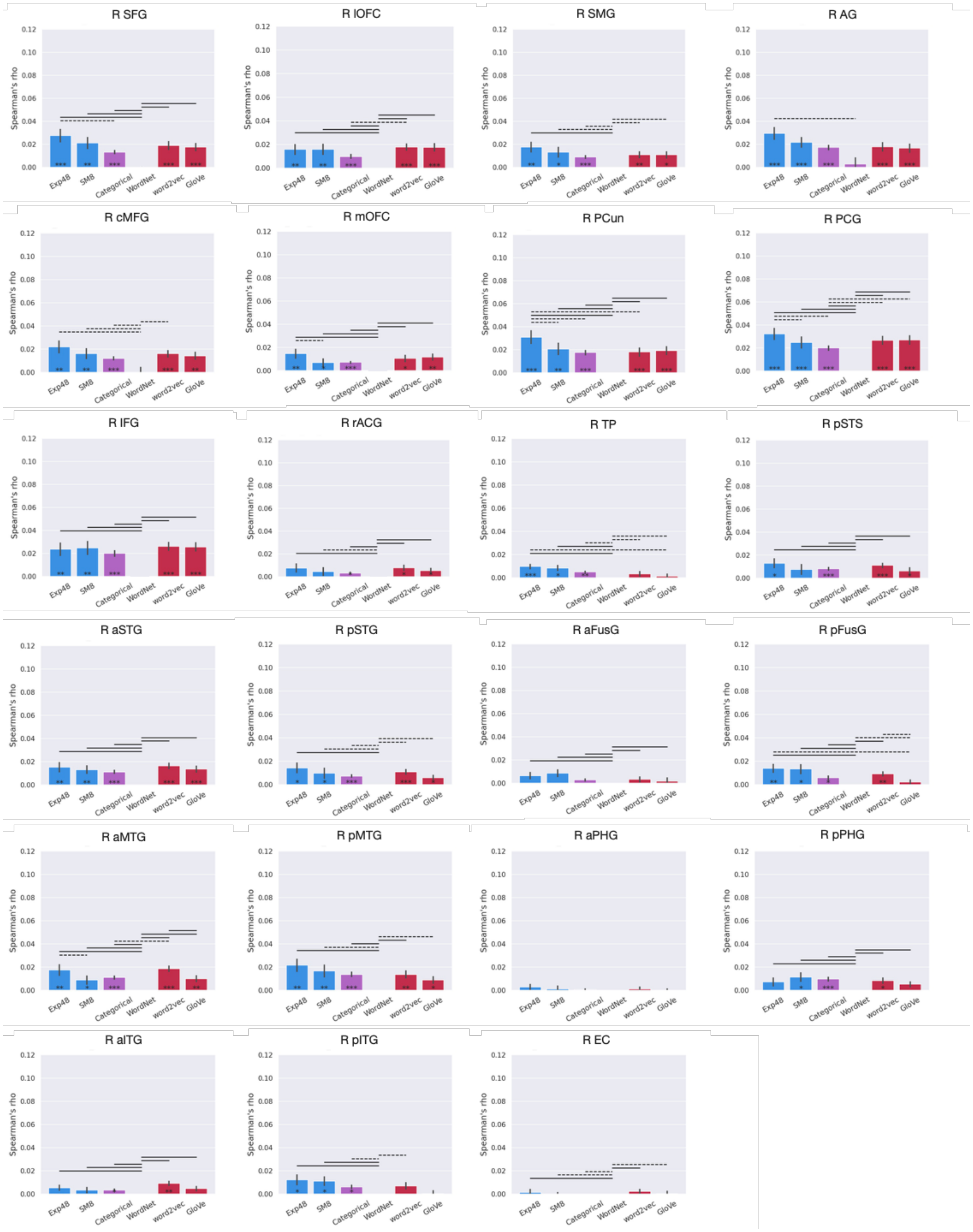


**Figure S10.** RSA results (across participants) for event concepts (right hemisphere anatomical ROIs). Color and symbol conventions as in Figure 3.

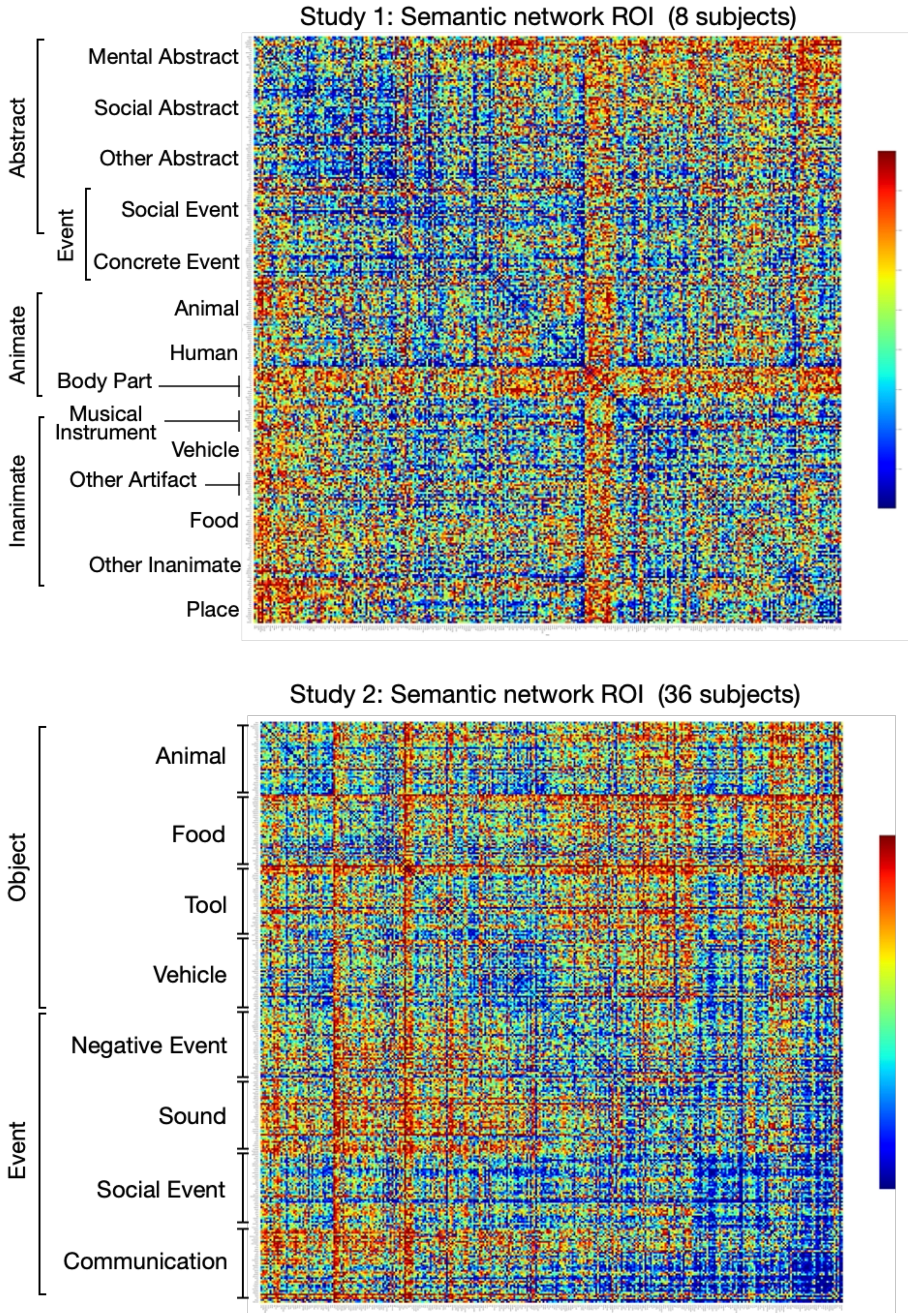


**Figure S11.** Neural RDMs, averaged across participants, for the semantic network ROI in Study 1 (top) and Study 2 (bottom).

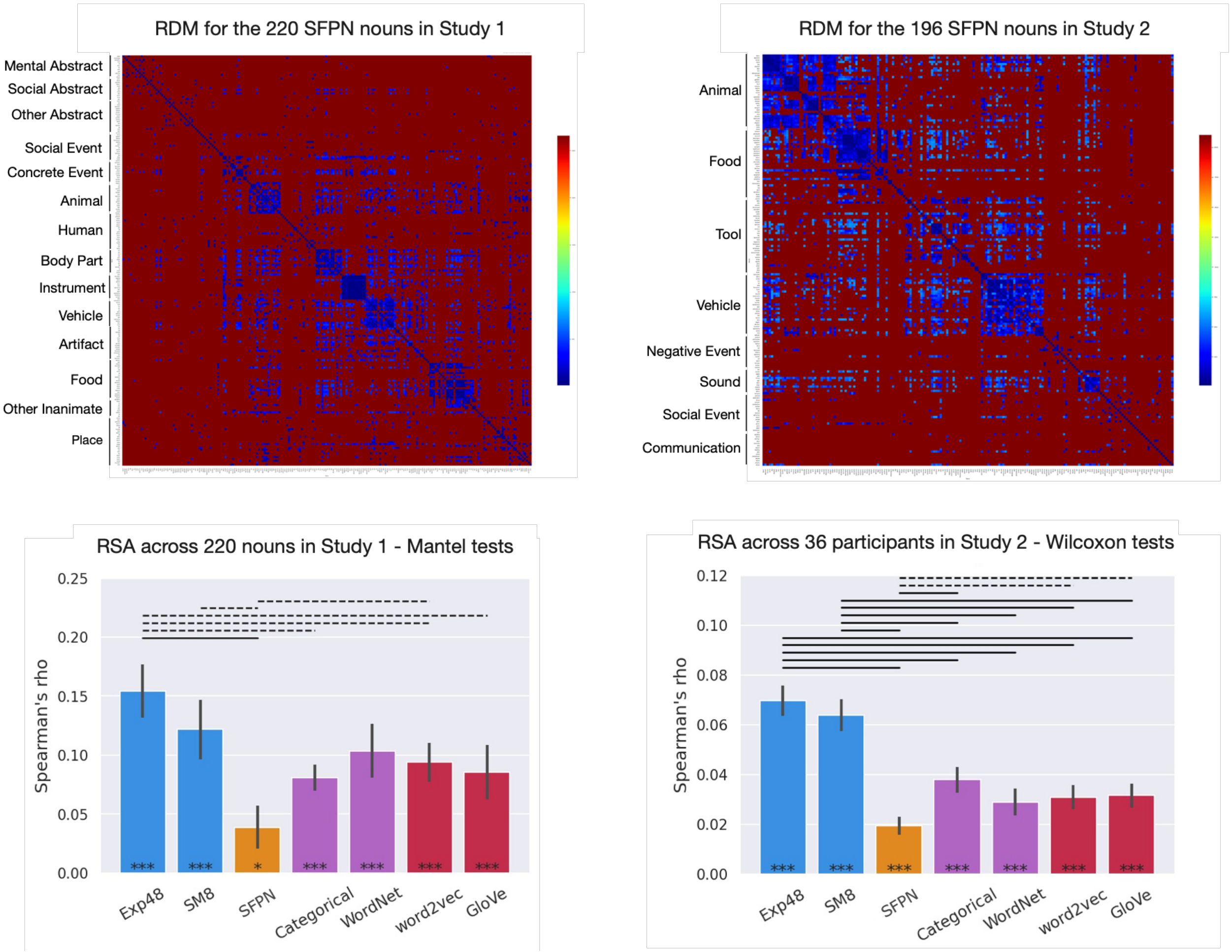


**Figure S12**. Top: RDMs for the SFPN models. Bottom: RSA results for the subset of nouns in each study for which SFPN norms are available.

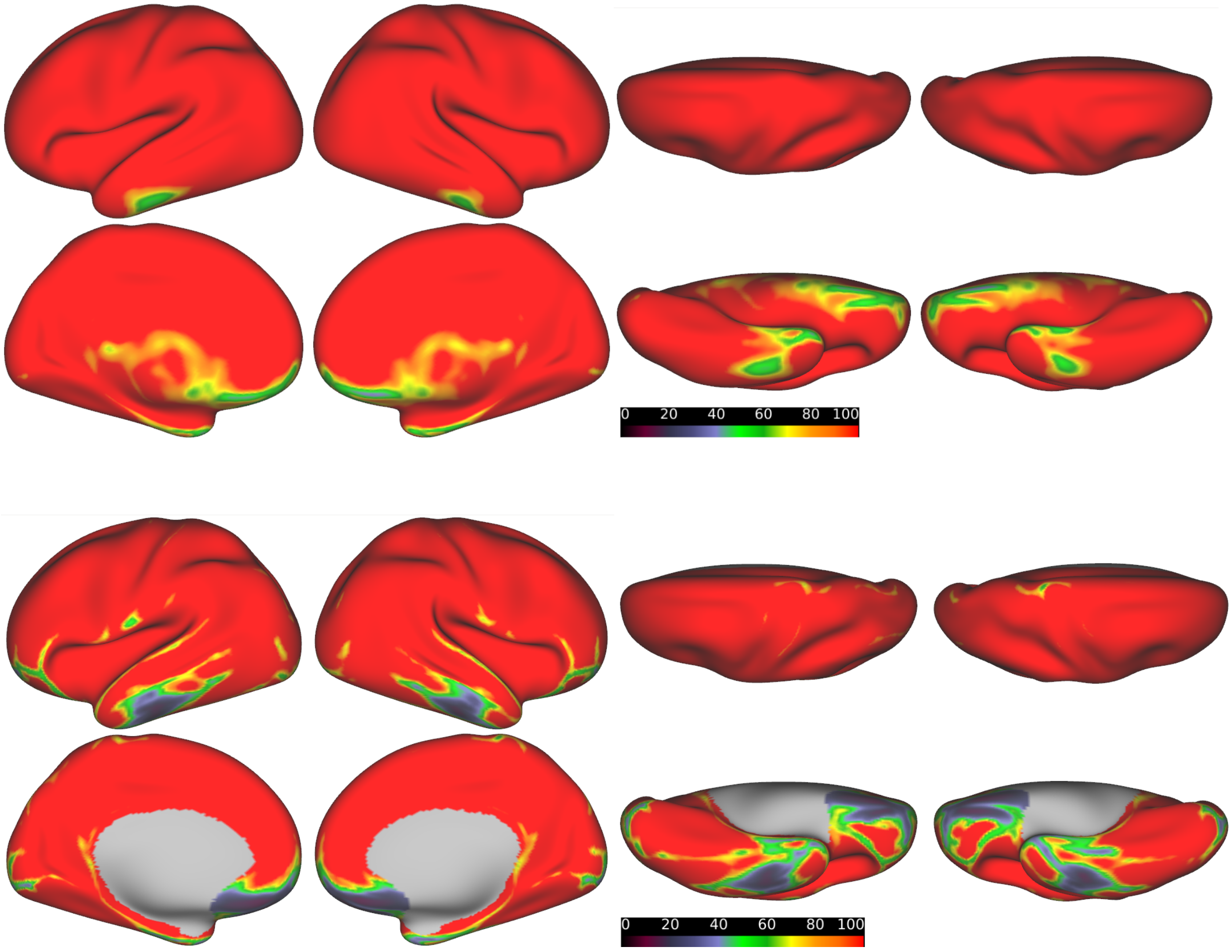


**Figure S13.** Temporal SNR maps for Study 1 (top) and Study 2 (bottom).

**Table S1.** Correlations between model-based RDMs in the two studies.

**Study 1**

| | Exp48 | SM8 | Categorical | WordNet | GloVe |
|---|---|---|---|---|---|
| SM8 | 0.59 | | | | |
| Categorical | 0.44 | 0.18 | | | |
| WordNet | 0.50 | 0.25 | 0.43 | | |
| GloVe | 0.43 | 0.32 | 0.23 | 0.25 | |
| word2vec | 0.37 | 0.25 | 0.22 | 0.23 | 0.60 |

**Study 2**

| | Exp48 | SM8 | Categorical | WordNet | GloVe |
|---|---|---|---|---|---|
| SM8 | 0.77 | | | | |
| Categorical | 0.73 | 0.46 | | | |
| WordNet | 0.51 | 0.30 | 0.68 | | |
| GloVe | 0.45 | 0.35 | 0.37 | 0.26 | |
| word2vec | 0.42 | 0.30 | 0.39 | 0.29 | 0.68 |

**Table S2.** Dimensions included in the Exp48 model.

| Dimension | Rating query<br>(to what degree do you think of this thing as…) | High Score Examples |
|---|---|---|
| Vision | being something you can easily see | moon, locomotive |
| Bright | being visually light or bright | sun, lightning |
| Dark | being visually dark | night, crow |
| Color | having a characteristic or defining color | grass, banana |
| Pattern | having a characteristic visual texture or pattern | tiger, pineapple |
| Motion | having a lot of visually observable movement | tornado, parade |
| Fast | having visible movement that is fast | rocket, cheetah |
| Slow | having visible movement that is slow | snail, cloud |
| Shape | having a characteristic visual shape or form | giraffe, spoon |
| Large | being large in size | volcano, ferry |
| Small | being small in size | bacterium, pea |
| Touch | easily recognizable by touch | toothbrush, sandpaper |
| Temperature | being either hot or cold to the touch | bonfire, ice |
| Texture | having either a smooth or rough texture to the touch | silk, stubble |
| Weight | being either light or heavy in weight | balloon, anvil |
| Pain | being associated with pain or physical discomfort | headache, bombing |
| Taste | having a characteristic taste | lemon, chocolate |
| Smell | having a characteristic smell | tobacco, barbecue |
| Mouth Action | associated with actions of the mouth | whistle, jaw |
| Hand Action | associated with actions of the arm, hand or fingers | keyboard, scissors |
| Foot Action | associated with actions of the legs or feet | soccer, bicycle |
| Manipulation | a physical object you have personally manipulated | fork, computer |
| Landmark | having a fixed location, as on a map | airport, library |
| Near | being near you, within reaching distance | foot, chair |
| Scene | belonging to a particular setting or physical location | oven, beach |
| Sound | having a characteristic or recognizable sound | rooster, piano |
| Audition | being something that you can easily hear | siren, thunder |
| Loud | making a loud sound | explosion, megaphone |
| Low pitch | having a low-pitched sound | tuba, growling |
| High pitch | having a high-pitched sound | whistle, dolphin |
| Path | showing motion along a particular direction or path | rocket, trolley |
| Toward | being associated with movement toward you | food, embrace |
| Away | being associated with movement away from you | cough, kite |
| Time | an event that occurs at a typical or predictable time | lunch, night |
| Duration | having a predictable duration, whether short or long | movie, year |
| Long Duration | an event that lasts for a long period of time | life, infinity |
| Short Duration | an event that lasts for a short period of time | sneeze, gunshot |
| Caused | caused by a preceding event, action, or situation | spill, honeymoon |
| Consequential | likely to have consequences | invasion, earthquake |
| Intention | having human-like intentions, plans, or goals | lobbyist, activist |
| Benefit | something that could help or benefit you or others | cure, peace |
| Harm | something that could cause harm to you or others | epidemic, wildfire |

| | | |
|---|---|---|
| Drive | something that motivates you to do something | duty, hope |
| Needs | something that would be hard to live without | shelter, water |
| Attention | someone or something that grabs your attention | scream, lightning |
| Pleasant | something that you find pleasant | vacation, cake |
| Unpleasant | something that you find unpleasant | pain, bombing |
| Arousal | something that makes you feel alert or excited | rollercoaster, lust |

**Table S3.** Lexical properties of the nouns included in Study 1. ON = orthographic neighborhood size. PN = phonological neighborhood size. Log frequency, ON, and PN are based on the HAL corpus. All data were compiled by the English Lexicon Project (8) (https://elexicon.wustl.edu/).

| | Letters | Phonemes | Syllables | Log Frequency | ON | PN | Age of Acquisition | Concreteness |
|---|---|---|---|---|---|---|---|---|
| Minimum | 3 | 1 | 1 | 4.0 | 0 | 0 | 2.7 | 1.2 |
| Maximum | 11.0 | 9.0 | 5.0 | 12.7 | 19 | 40 | 13.8 | 5.0 |
| Mean | 5.9 | 4.9 | 1.9 | 8.7 | 3.6 | 7.6 | 6.7 | 4.1 |

**Table S4.** Noun stimuli included in Study 1.

| Word | Category | Word | Category | Word | Category |
|---|---|---|---|---|---|
| belief | mental | sin | social abstract | battle | social event |
| hope | mental | snub | social abstract | carnival | social event |
| intellect | mental | testimony | social abstract | circus | social event |
| knowledge | mental | treaty | social abstract | debate | social event |
| optimism | mental | tribute | social abstract | speech | social event |
| sympathy | mental | truce | social abstract | election | social event |
| trust | mental | trial | social abstract | festival | social event |
| wit | mental | attribute | abstract | funeral | social event |
| animosity | mental | year | abstract | honeymoon | social event |
| awe | mental | curse | abstract | matinee | social event |
| delirium | mental | worth | abstract | meeting | social event |
| dread | mental | day | abstract | oration | social event |
| envy | mental | fate | abstract | party | social event |
| fun | mental | fee | abstract | bonfire | social event |
| gratitude | mental | folly | abstract | rally | social event |
| grief | mental | era | abstract | vacation | social event |
| guilt | mental | heredity | abstract | musical | social event |
| ire | mental | home | abstract | soccer | social event |
| jealousy | mental | semester | abstract | riot | social event |
| joy | mental | hygiene | abstract | parade | social event |
| love | mental | infinity | abstract | applause | social event |
| malice | mental | majority | abstract | dinner | social event |
| shame | mental | number | abstract | embrace | concrete event |
| torment | mental | morning | abstract | handshake | concrete event |
| woe | mental | peace | abstract | kiss | concrete event |
| victim | social abstract | problem | abstract | avalanche | concrete event |
| bribe | social abstract | quantity | abstract | belch | concrete event |
| deceit | social abstract | evening | abstract | clang | concrete event |
| etiquette | social abstract | reality | abstract | cough | concrete event |
| fallacy | social abstract | accident | abstract | cyclone | concrete event |
| grievance | social abstract | role | abstract | downpour | concrete event |
| insult | social abstract | patent | abstract | explosion | concrete event |
| joke | social abstract | winter | abstract | fireworks | concrete event |
| loan | social abstract | sum | abstract | flood | concrete event |
| mercy | social abstract | summer | abstract | gasp | concrete event |
| moral | social abstract | tax | abstract | gunshot | concrete event |
| perjury | social abstract | truth | abstract | hailstorm | concrete event |
| plea | social abstract | vice | abstract | hurricane | concrete event |
| rumor | social abstract | night | abstract | landslide | concrete event |

| Word | Category | Word | Category | Word | Category |
|---|---|---|---|---|---|
| lightning | concrete event | girl | human | saxophone | instrument |
| ricochet | concrete event | guard | human | trombone | instrument |
| whine | concrete event | farmer | human | trumpet | instrument |
| scream | concrete event | voter | human | tuba | instrument |
| screech | concrete event | doctor | human | ambulance | vehicle |
| squeal | concrete event | driver | human | bicycle | vehicle |
| stampede | concrete event | worker | human | boat | vehicle |
| storm | concrete event | army | human group | bus | vehicle |
| thunder | concrete event | audience | human group | cab | vehicle |
| tornado | concrete event | choir | human group | car | vehicle |
| alligator | animal | couple | human group | carriage | vehicle |
| ant | animal | jury | human group | plane | vehicle |
| bee | animal | mob | human group | sailboat | vehicle |
| butterfly | animal | family | human group | scooter | vehicle |
| camel | animal | arm | body part | sled | vehicle |
| cheetah | animal | eye | body part | submarine | vehicle |
| crow | animal | foot | body part | subway | vehicle |
| elephant | animal | hair | body part | train | vehicle |
| turtle | animal | hand | body part | truck | vehicle |
| whale | animal | finger | body part | van | vehicle |
| goldfish | animal | toe | body part | axe | artifact |
| hawk | animal | jaw | body part | ball | artifact |
| snake | animal | leg | body part | baseball | artifact |
| tiger | animal | shoulder | body part | bed | artifact |
| monkey | animal | lip | body part | bell | artifact |
| moose | animal | mouth | body part | comb | artifact |
| mosquito | animal | muscle | body part | dime | artifact |
| penguin | animal | nose | body part | door | artifact |
| boy | human | accordion | instrument | elevator | artifact |
| criminal | human | bagpipe | instrument | escalator | artifact |
| businessman | human | chime | instrument | fountain | artifact |
| politician | human | clarinet | instrument | football | artifact |
| soldier | human | drum | instrument | limousine | artifact |
| student | human | flute | instrument | newspaper | artifact |
| terrorist | human | gong | instrument | pan | artifact |
| tourist | human | harmonica | instrument | rocket | artifact |
| parent | human | harp | instrument | television | artifact |
| patient | human | mandolin | instrument | tobacco | artifact |
| man | human | piano | instrument | window | artifact |

| Word | Category | Word | Category |
|---|---|---|---|
| beer | food | feather | inanimate |
| cheese | food | ice | inanimate |
| chocolate | food | sun | inanimate |
| corn | food | school | manmade place |
| egg | food | store | manmade place |
| honey | food | theater | manmade place |
| jam | food | cafeteria | manmade place |
| lemonade | food | cathedral | manmade place |
| mustard | food | church | manmade place |
| pie | food | college | manmade place |
| rum | food | hall | manmade place |
| spaghetti | food | hospital | manmade place |
| tea | food | hotel | manmade place |
| apricot | food | kitchen | manmade place |
| banana | food | lab | manmade place |
| blueberry | food | office | manmade place |
| carrot | food | prison | manmade place |
| cherry | food | airport | manmade place |
| chestnut | food | street | manmade place |
| coffee | food | bridge | manmade place |
| eggplant | food | garden | manmade place |
| plum | food | park | manmade place |
| raspberry | food | zoo | manmade place |
| tangerine | food | farm | manmade place |
| tomato | food | bay | natural place |
| dandelion | inanimate | beach | natural place |
| elm | inanimate | forest | natural place |
| flower | inanimate | island | natural place |
| ivy | inanimate | jungle | natural place |
| oak | inanimate | mountain | natural place |
| rose | inanimate | prairie | natural place |
| tulip | inanimate | river | natural place |
| cloud | inanimate | volcano | natural place |

**Table S5.** Lexical properties of the nouns included in Study 2.

| | Letters | Phonemes | Syllables | Log Frequency | ON. | PN | Age of Acquisition | Concreteness |
|---|---|---|---|---|---|---|---|---|
| Minimum | 3 | 2 | 1 | 2.94 | 0 | 0 | 2.7 | 2.1 |
| Maximum | 13 | 11 | 5 | 12.44 | 18 | 41 | 14.8 | 5.0 |
| Mean | 6.9 | 5.7 | 2.1 | 7.61 | 2.2 | 5.1 | 7.2 | 4.2 |
| Mean Object | 6.7 | 5.5 | 2.1 | 7.66 | 2.4 | 5.7 | 6.0 | 4.9 |
| Mean Event | 7.2 | 5.8 | 2.2 | 7.57 | 1.9 | 4.5 | 8.4 | 3.6 |

**Table S6.** Noun stimuli included in Study 2.

| Word | Category | Word | Category | Word | Category |
|---|---|---|---|---|---|
| alligator | animal | whale | animal | tobacco | food |
| ant | animal | asparagus | food | tomato | food |
| baboon | animal | banana | food | anchor | tool |
| bison | animal | bean | food | axe | tool |
| butterfly | animal | beer | food | baseball | tool |
| cardinal | animal | blueberry | food | binoculars | tool |
| caterpillar | animal | bread | food | book | tool |
| chameleon | animal | broccoli | food | calculator | tool |
| cheetah | animal | carrot | food | camera | tool |
| chicken | animal | champagne | food | candle | tool |
| chimpanzee | animal | cheese | food | cash | tool |
| chipmunk | animal | cherry | food | comb | tool |
| cricket | animal | chestnut | food | corkscrew | tool |
| crow | animal | chocolate | food | crutches | tool |
| dog | animal | cider | food | dime | tool |
| dolphin | animal | coffee | food | faucet | tool |
| duck | animal | cranberry | food | football | tool |
| elephant | animal | cucumber | food | fork | tool |
| fish | animal | custard | food | glass | tool |
| goldfish | animal | dandelion | food | hairbrush | tool |
| hamster | animal | egg | food | hammer | tool |
| hawk | animal | eggplant | food | handsaw | tool |
| hippopotamus | animal | flower | food | hoe | tool |
| horse | animal | ham | food | key | tool |
| jackal | animal | honey | food | keyboard | tool |
| lion | animal | jam | food | ladle | tool |
| monkey | animal | ketchup | food | magazine | tool |
| moose | animal | lemonade | food | microscope | tool |
| mosquito | animal | milk | food | newspaper | tool |
| mouse | animal | mushroom | food | pencil | tool |
| octopus | animal | mustard | food | rake | tool |
| penguin | animal | nectarine | food | sandpaper | tool |
| rhinoceros | animal | pineapple | food | scissors | tool |
| salmon | animal | plant | food | skillet | tool |
| snake | animal | pudding | food | spatula | tool |
| tiger | animal | pumpkin | food | stapler | tool |
| trout | animal | raspberry | food | stethoscope | tool |
| turkey | animal | sauerkraut | food | straw | tool |
| turtle | animal | spaghetti | food | thermometer | tool |

| Word | Category | Word | Category | Word | Category |
|---|---|---|---|---|---|
| ticket | tool | trolley | vehicle | twister | negative event |
| tongs | tool | truck | vehicle | volcano | negative event |
| umbrella | tool | van | vehicle | war | negative event |
| ambulance | vehicle | wagon | vehicle | whirlwind | negative event |
| automobile | vehicle | avalanche | negative event | wildfire | negative event |
| barge | vehicle | battle | negative event | applause | sound |
| bicycle | vehicle | blizzard | negative event | bang | sound |
| boat | vehicle | bombing | negative event | bellowing | sound |
| bobsled | vehicle | brawl | negative event | boom | sound |
| bus | vehicle | cyclone | negative event | chattering | sound |
| canoe | vehicle | downpour | negative event | chuckle | sound |
| car | vehicle | drought | negative event | clapping | sound |
| carriage | vehicle | earthquake | negative event | clattering | sound |
| convertible | vehicle | epidemic | negative event | crackle | sound |
| elevator | vehicle | explosion | negative event | crescendo | sound |
| escalator | vehicle | famine | negative event | giggle | sound |
| ferry | vehicle | flood | negative event | groaning | sound |
| glider | vehicle | gunshot | negative event | growling | sound |
| helicopter | vehicle | gust | negative event | grunt | sound |
| jeep | vehicle | hail | negative event | gulp | sound |
| limousine | vehicle | hailstorm | negative event | hiccup | sound |
| locomotive | vehicle | hurricane | negative event | jingle | sound |
| motorcycle | vehicle | inferno | negative event | laughter | sound |
| plane | vehicle | invasion | negative event | melody | sound |
| rocket | vehicle | landslide | negative event | murmuring | sound |
| rowboat | vehicle | lightning | negative event | reverberation | sound |
| sailboat | vehicle | monsoon | negative event | roaring | sound |
| scooter | vehicle | murder | negative event | rumble | sound |
| skateboard | vehicle | outbreak | negative event | rustle | sound |
| sled | vehicle | plague | negative event | screaming | sound |
| sleigh | vehicle | raid | negative event | screeching | sound |
| steamer | vehicle | riot | negative event | shrieking | sound |
| streetcar | vehicle | shooting | negative event | sigh | sound |
| submarine | vehicle | squall | negative event | siren | sound |
| subway | vehicle | stampede | negative event | sizzle | sound |
| taxi | vehicle | storm | negative event | snap | sound |
| tractor | vehicle | tempest | negative event | sneeze | sound |
| train | vehicle | thunderstorm | negative event | sobbing | sound |
| tricycle | vehicle | tornado | negative event | squeaking | sound |

| Word | Category | Word | Category | Word | Category |
|---|---|---|---|---|---|
| squeal | sound | luncheon | social event | discourse | communication |
| thumping | sound | march | social event | dispute | communication |
| thunderclap | sound | musical | social event | eulogy | communication |
| wheezing | sound | outing | social event | greeting | communication |
| whimpering | sound | pageant | social event | grievance | communication |
| whine | sound | parade | social event | huddle | communication |
| banquet | social event | party | social event | interrogation | communication |
| bash | social event | picnic | social event | joke | communication |
| carnival | social event | prom | social event | lecture | communication |
| celebration | social event | rally | social event | lesson | communication |
| christening | social event | reception | social event | meeting | communication |
| circus | social event | reunion | social event | plea | communication |
| cocktails | social event | safari | social event | praise | communication |
| concert | social event | symphony | social event | protest | communication |
| conference | social event | tour | social event | quarrel | communication |
| contest | social event | tournament | social event | rant | communication |
| convention | social event | wedding | social event | rebuke | communication |
| cookout | social event | advice | communication | rebuttal | communication |
| cruise | social event | apology | communication | recitation | communication |
| dance | social event | class | communication | sermon | communication |
| expedition | social event | commemoration | communication | showdown | communication |
| expo | social event | comment | communication | squabble | communication |
| fair | social event | commentary | communication | testimony | communication |
| feast | social event | complaint | communication | thanks | communication |
| festival | social event | compliment | communication | threat | communication |
| fiesta | social event | debate | communication | trial | communication |
| gathering | social event | denial | communication | tribute | communication |
| housewarming | social event | deposition | communication | wisecrack | communication |
| jubilee | social event | dictation | communication | | |

**Table S7.** Mean group-averaged tSNR for each anatomically defined ROI.

| ROI | Study 1 | Study 2 |
|---|---|---|
| L AG | 151 | 125 |
| R AG | 154 | 138 |
| L SMG | 157 | 129 |
| R SMG | 165 | 143 |
| L PCun | 141 | 125 |
| R PCun | 148 | 134 |
| L PCG | 126 | 121 |
| R PCG | 131 | 121 |
| L lat OFC | 95 | 70 |
| R lat OFC | 93 | 74 |
| L mOFC | 59 | 33 |
| R mOFC | 63 | 38 |
| L SFG | 153 | 133 |
| R SFG | 153 | 128 |
| L cMFG | 146 | 130 |
| R cMFG | 149 | 132 |
| L IFG | 137 | 109 |
| R IFG | 128 | 117 |
| L rAC | 124 | 102 |
| R rAC | 136 | 106 |
| L pSTS | 136 | 126 |
| R pSTS | 179 | 159 |
| L aSTG | 134 | 102 |
| R aSTG | 138 | 109 |
| L pSTG | 172 | 108 |
| R pSTG | 177 | 123 |
| L aMTG | 150 | 70 |
| R aMTG | 149 | 78 |
| L pMTG | 174 | 86 |
| R pMTG | 163 | 97 |
| L aITG | 106 | 34 |
| R aITG | 111 | 36 |
| L pITG | 137 | 67 |
| R pITG | 127 | 69 |
| L aFusG | 87 | 29 |
| R aFusG | 82 | 33 |
| L pFusG | 148 | 106 |
| R pFusG | 137 | 106 |
| L aPHG | 98 | 42 |

| | | |
|---|---|---|
| R aPHG | 86 | 37 |
| L pPHG | 107 | 93 |
| R pPHG | 105 | 93 |
| L EC | 81 | 42 |
| R EC | 80 | 28 |
| L TP | 80 | 53 |
| R TP | 81 | 46 |
